# Jumonji-C domain containing demethylase 1 is critical for heterochromatin organization in *Plasmodium falciparum*

**DOI:** 10.1101/2025.10.08.680194

**Authors:** Jonas Gockel, Matthias Wyss, Parul Singh, Abhishek Kanyal, Dominique Keller, Travis Basson, Beatriz Graça, Teja Rus, Jessica Bryant, Till Voss, Richárd Bártfai

**Affiliations:** Radboud University, Department of Molecular Biology, Nijmegen, the Netherlands; Swiss Tropical and Public Health Institute, Department of Medical Parasitology and Infection Biology, Basel, Switzerland; University of Basel, Basel, Switzerland; Institut Pasteur, Université Paris Cité, INSERM U1201, CNRS EMR9195, Biology of Host-Parasite Interactions Unit, Paris, France

## Abstract

Despite the central importance of epigenetic regulation in enabling adaptation and pathogenicity of malaria parasites, mechanisms controlling heterochromatin distribution in *Plasmodium falciparum* are poorly understood. Here, we identified *P. falciparum* Jumonji C domain-containing histone demethylase 1, PfJmjC1, as a key regulator of heterochromatin. Parasites lacking PfJmjC1 are viable *in vitro* but display slightly reduced multiplication rates compared to wild type parasites. Interestingly, PfJmjC1 depletion leads to major heterochromatin reorganization, involving *de novo* heterochromatin formation over non-essential, GC-rich euchromatic genes and reduction of heterochromatin at some chromosome ends, eventually altering the 3D genome architecture. Importantly, this heterochromatin reorganization results in the deregulation of cell surface antigen expression and failure of parasites to induce gametocyte production in response to nutrient deprivation. Collectively, our findings highlight the crucial role of PfJmjC1 in regulating heterochromatin distribution and life cycle progression in this deadly pathogen.

## Introduction

Malaria remains a devastating global health challenge, causing around 260 million infections resulting in approximately 600.000 deaths each year^1^. The disease is caused by intracellular parasites of the *Plasmodium* genus, which invade human red blood cells and multiply excessively. Malaria eradication efforts are compromised by high phenotypic variability between and even within parasite strains, facilitating immune escape, chronic infection^2,3^ and drug resistance^4^. Clonal phenotypic variability is primarily enabled by epigenetic mechanisms^5,6^, and its major driver is variation in organization of transcriptionally-repressive heterochromatin^7,8^. Heterochromatin in *Plasmodia* is characterized by trimethylation on histone 3 lysine 9 (H3K9me3) and stands in opposition to transcriptionally-permissive euchromatin, which is generally marked by acetylation of various histone tails^9^.

In *Plasmodium falciparum*, heterochromatin is distinctively deposited in the subtelomeric regions at all chromosome ends, but also at some intrachromosomal islands^8^. The genes repressed by these heterochromatic domains are mainly members of multigene families such as *var*, *stevor* and *rifins* that are involved in host cell remodeling and antigenic variation^10–13^. Importantly, variation in heterochromatin organization between individual parasites leads to expression of distinct subsets of these gene family members and hence enables clonal phenotypic variation and adaptation via a “bet-hedging” strategy^2,14^. In addition to multigene families, genes involved in life cycle progression such as *pfap2-g*, encoding the master regulator of gametocytogenesis, are also regulated by heterochromatin^7^. Upstream of *pfap2-g*, gametocyte development protein 1 (GDV1) negatively impacts heterochromatin occupancy over the *pfap2-g* locus, leading to activation of *pfap2-g* expression and thereby initiation of gametocytogenesis^15^. During sexual differentiation, heterochromatin is not only removed from the *pfap2-g* locus and some gametocyte-specific genes but also expands in some subtelomeric regions to silence asexual stage-specific genes such as knob-associated histidine-rich protein (*kahrp*)^8^. Accordingly, proper control of heterochromatin occupancy is imperative for adaptation to environmental challenges and life cycle progression of malaria parasites, and it is crucial to understand how dynamic rearrangements of heterochromatin are orchestrated.

During heterochromatin formation and maintenance, H3K9me3 is deposited by histone methyltransferases (HMT) and in turn bound by heterochromatin protein 1 (HP1)^16^. These HMTs bind to heterochromatin and methylate neighboring histone tails, thus enabling heterochromatin spreading^17^. This self-propagating characteristic leads to the formation of broad heterochromatic regions and accompanying gene repression. In *P. falciparum*, several factors such as the histone deacetylases PfSir2a/PfSir2b^5,18^ and PfHDA2^19^, the PfHP1 antagonist GDV1^15^ , the HMT PfSET2^20^ and others have been shown to affect heterochromatin. PfSET3 has been postulated as the HMT that deposits the H3K9me3 mark^6,21^. However, the mechanisms by which these proteins cooperate to facilitate heterochromatin formation and spread are not yet understood.

As heterochromatin has the intrinsic ability to spread to neighboring regions, it must be tightly regulated to ensure proper gene expression as well as heterochromatin occupancy. In model eukaryotes, several mechanisms of limiting heterochromatin spread (so called heterochromatin boundaries) have been proposed^22^, such as limited availability of heterochromatin constituents^23^, steric hinderance^24–26^ or active gene expression^27,28^. Another possible mechanism is the active removal of deposited H3K9me3 marks by histone demethylases (HDMs), which directly antagonize HMT activity^29^. Currently, while heterochromatin assembly and maintenance in *P. falciparum* has been the subject of several studies^8,14,30,31^, the mechanism controlling heterochromatin spread in these parasites is not understood.

While histone demethylases are prime candidates that could be involved in heterochromatin boundary maintenance, their functions in this regard have not been investigated in *P. falciparum*. Two classes of demethylases are present in malaria parasites: Jumonji-C domain containing demethylases (JmjCs) and lysine-specific demethylases (LSDs)^21^. LSDs in general are not able to remove H3K9 tri-methylation due to the limited size of their enzymatic active pocket and the lack of protonated lysine in trimethyl residues^32–34^. However, LSDs can still demethylate H3K9me2 and H3K9me1^32,35–37^. Conversely, as JmjCs remove methylation by oxidation, they are able to demethylate tri-,di and mono-methylated states of H3K9^32^. Intriguingly, most demethylases encoded in the *P. falciparum* genome are dispensable for asexual growth in blood stage parasites *in vitro*. PfLSD1, PfJmjC1 and PfJmjC2 null mutants were shown to be viable^20^. While PfLSD2 is also dispensable for asexual growth^38^, it is expressed in and likely required for very early gametocytes^39–41^. PfJmjC3 was also shown to be non-essential in asexual blood stages in a transposon-mediated gene essentiality screen^38^. Small molecule inhibitors targeting recombinant PfJmjC3 have been shown to disrupt *P. falciparum* development^42^. This effect, however, might be due to limited specificity of the compound and broader inhibition of demethylase activities in the parasite and could potentially point to some redundance between demethylases. Of all predicted demethylases in *P. falciparum*, PfJmjC1 appears to be the most likely candidate involved in controlling heterochromatin as native co-immunoprecipitation (Co-IP) experiments suggest PfJmjC1 interacts with PfHP1^15^, and potentially in antagonizing relevant HMT activities.

In this study, we show that PfJmjC1 does indeed interact with heterochromatin and associated proteins. Importantly, we also demonstrate that PfJmjC1 is essential for maintaining proper heterochromatin distribution. PfJmjC1 knockout parasites display substantially reorganized heterochromatin including both *de novo* heterochromatin formation in euchromatic regions as well as reduction of heterochromatin occupancy at chromosome ends, while the overall spread of existing heterochromatin domains is unaffected. This re-distribution of heterochromatin occurs between regions with similar GC-content and affects weakly-expressed euchromatic genes that are not essential in asexual parasites. Furthermore, a subset of genes that gain heterochromatin upon PfJmjC1 depletion show increased long-range interactions with heterochromatic multigene families, indicating substantial rearrangements in nuclear architecture. Notably, PfJmjC1 knockout parasites lose their capacity to trigger gametocyte production under nutrient-poor conditions, underlining the critical role for PfJmjC1 in parasite life cycle progression and transmission.

## Results

### PfJmjC1 interacts with heterochromatin and associated proteins

To test the hypothesis that antagonizing activity of HDMs towards HMTs limits heterochromatin spread and assists proper heterochromatin maintenance (Fig. 1a), we set out to functionally characterize the Jumonji(Jmj)-C domain containing demethylase PfJmjC1. This protein (unlike PfJmjC2/3 and PfLSD1/2,) is predicted to contain a Jumonji-N and a H5HC2 type zinc finger (H5HC2-ZF) domain in addition to the JmjC domain (Fig. 1b, Extended Data Fig. 1a)^43^. Orthologs of PfJmjC1 are also present in other *Plasmodium* species such as *P. vivax*, showing ∼40% identity in pairwise alignment with PfJmjC1 mainly in the conserved JmjN, JmjC and H5HC2-ZF domains (Extended Data Fig. 1b). Multi-sequence alignment further highlights conservation of the binding sites for both JmjC cofactors α-ketoglutarate and Fe-(II) ions within the JmjC domain of human and yeast demethylases with similar domain architecture (Fig. 1c), indicating that PfJmjC1 is likely an active histone demethylase enzyme.

**Fig. 1:**
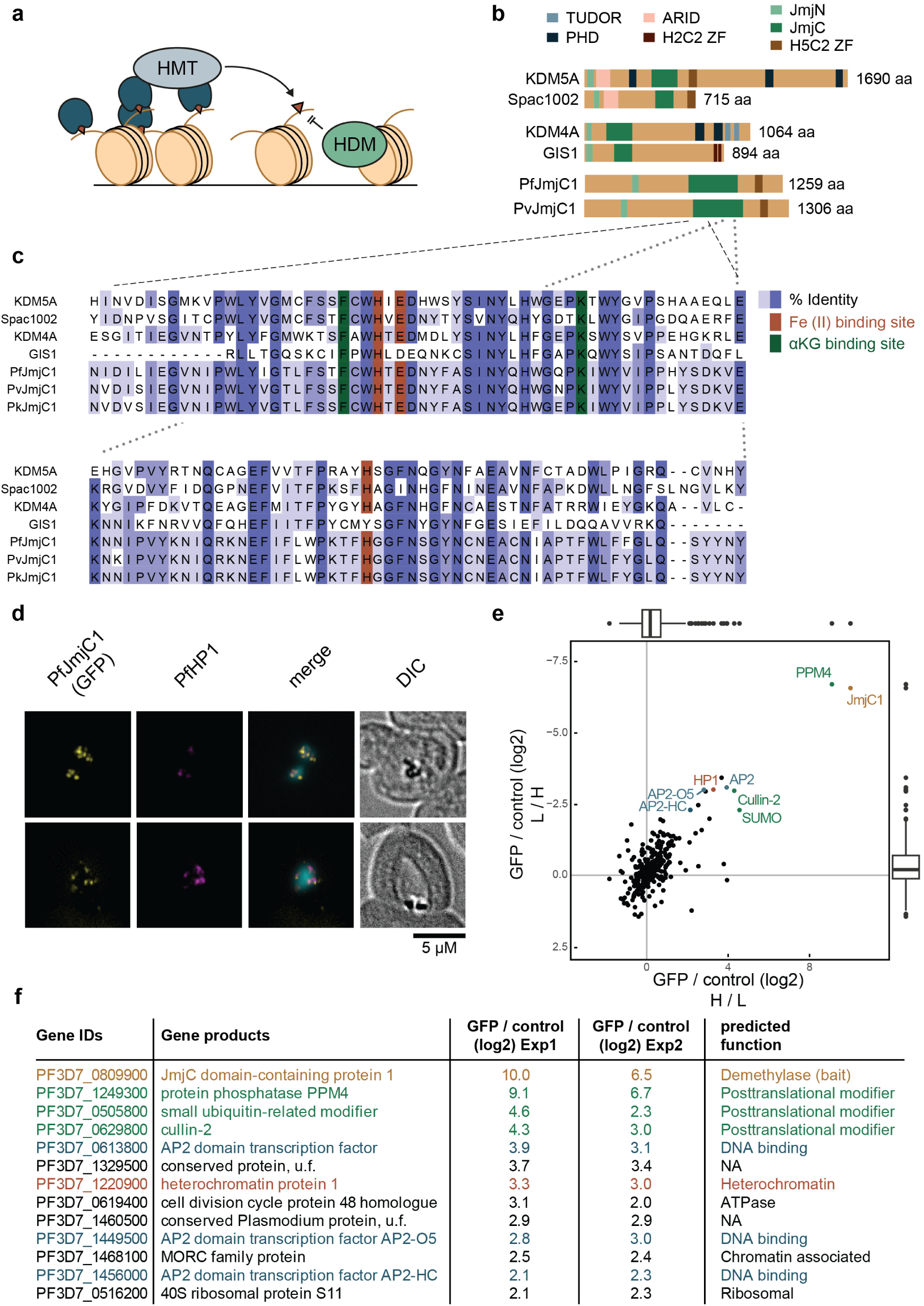
JmjC1 interacts with heterochromatin. **a)** Proposed interaction of histone methyltransferases (HMTs) and Histone demethylases (HDMs) at heterochromatin – euchromatin boundaries. Removal of H3K9me3 by demethylase enzymes could antagonize spread of heterochromatin. **b)** Domain structure of different JmjN/JmjC domain containing proteins in Homo sapiens (KDM5A/KDM4A), Schizosaccharomyces pombe (Spac1002), Saccharomyces cerevisiae (GIS1) and P. falciparum/P. vivax JmjC1, as annotated by InterProtScan. **c)** Multiple sequence alignment of the JmjC domains of proteins shown in panel b, highlighting two separate parts of the JmjC domains with co-factor binding sites. Fe(II) binding sites are marked in red, alpha-ketoglutarate (αKG) binding sites in green and level of amino acid (aa) identity with a blue hue. The upper and lower alignment displays regions corresponding to PfJmjC1 aa 681 – 738, and aa 917 – 972, respectively. **d)** Two representative examples of IFAs of PfJmjC1-GFP and PfHP1 in the 3D7/PfJmjC1-GFP parasite line. Anti-GFP signals are shown in yellow and anti-PfHP1 in magenta. DAPI was used to stain DNA (blue). **e)** Scatterplot of proteins enriched in the eluates of PfJmjC1-GFP Co-IP experiments from two separate experiments with swapped isotope labels (**L**ight and **H**eavy). Boxplots show quartiles 1 to 3, median is indicated as a thick line. **f)** Enriched proteins identified in the Co-IP experiments, with corresponding gene product names, anti-GFP/control enrichment values for both experiments and predicted functions based on information retrieved from PlasmoDB (www.plasmodb.org)^44^.

The postulated role for PfJmjC1 in heterochromatin maintenance would require at least a temporal association of PfJmjC1 to heterochromatin. Previous PfHP1 Co-IP data demonstrated a potential interaction with PfJmjC1 and did not identify any other demethylases^15^. To validate interactions of PfJmjC1 with heterochromatin, we constructed a parasite line expressing endogenous PfJmjC1 tagged with GFP at the C terminus (3D7/JmjC1-GFP) (Extended Data Figure 2a,b) and investigated co-localization of PfJmjC1-GFP with PfHP1 using immunofluorescence assay (IFA). PfJmjC1-GFP was most prominently detected at the trophozoite and early schizont stages, where it displayed nuclear localization with strong local enrichments at the nuclear periphery (Fig. 1d, Extended Data Fig. 2c). These perinuclear JmjC1-GFP foci showed substantial co-localization with PfHP1 foci, further supporting the association of PfJmjC1 with heterochromatin.

To identify interactors of PfJmjC1 potentially involved in aspects of (hetero)chromatin organization, we performed Co-IPs of PfJmjC1-GFP followed by mass spectrometry. Importantly, PfJmjC1 itself and PfHP1 were both clearly enriched in two independent experiments, showing that the pull-down was efficient and further confirming the interaction between PfJmjC1 and heterochromatin (Fig. 1e). Other interactors could be classified into two broad categories (Fig. 1f): i) ApiAP2 transcription factors with DNA-binding properties (PfAP2-HC, PfAP2-O5 and PF3D7_0613800) that might be involved in the recruitment of PfJmjC1 to chromatin; and ii) proteins involved in posttranslational modification (PfSUMO, PfPPM4, PfCullin-2), which could potentially synergize with PfJmjC1 in altering the chromatin structure. Out of these interactors, the phosphatase PfPPM4 exhibited the strongest enrichment, indicating stronger and/or more direct interaction with PfJmjC1.

Collectively, these experiments demonstrate spatial and physical association between PfJmjC1 and heterochromatin as well as heterochromatin-associated proteins that might collaborate in heterochromatin regulation.

### PfJmjC1 knockout leads to heterochromatin rearrangements

In order to test if PfJmjC1 has a function in heterochromatin maintenance/organization, we generated PfJmjC1 knockout (KO) parasites by replacing parts of the coding sequence with a human dihydrofolate reductase (h*dhfr*) drug resistance cassette (3D7/JmjC1-KO)(Extended Data Fig. 3a,c). As we cannot exclude the possibility that PfJmjC2 may also be involved in heterochromatin maintenance, we also generated a PfJmjC2 KO line (3D7/JmjC2-KO) by disrupting the gene via insertion of a blasticidin deaminase (*bsd*) resistance cassette (Extended Data Fig. 3b,d). 3D7/JmjC1-KO and 3D7/JmjC2-KO parasites were both easily obtained after drug selection of transfected parasites, confirming that both enzymes are dispensable for asexual blood stage development, as shown previously^20,38^. However, by quantifying parasite multiplication using flow cytometry we observed that 3D7/JmjC1-KO and 3D7/JmjC2-KO parasites multiplied at a slightly reduced rate (9.1 fold; ±0.4 s.d.; 8.2 fold, ±1.1 s.d.; respectively) compared to the 3D7 wild type parental strain (10.6 fold; ±0.7 s.d.) (Fig. 2a). Since PfJmjC2 could potentially compensate for the loss of PfJmjC1 function, and *vice versa*, we also disrupted the *pfjmjc2* locus in 3D7/JmjC1-KO parasites to generate a JmjC1/2 double loss-of-function mutant (3D7/JmjC1/2-dKO) (Fig. 2a, Extended Data Figure 3b,d). As shown in Fig. 2a, the 3D7/JmjC1/2-dKO double null mutant is also viable and displays a multiplication rate (7.7 fold; ±0.6 s.d.) that is slightly lower compared to the PfJmjC1 and PfJmjC2 single KO line. Based on microscopic inspection of parasite morphology and the multiplication assay data, neither the single nor the double KO lines display any obvious defects in stage progression throughout intra-erythrocytic schizogony. Hence, while the absence of PfJmjC1 and/or PfJmjC2 lead to slightly reduced growth rates, 3D7/JmjC1-KO, 3D7/JmjC2-KO and 3D7/JmjC1/2-dKO parasites are still able to undergo efficient asexual replication.

**Fig. 2:**
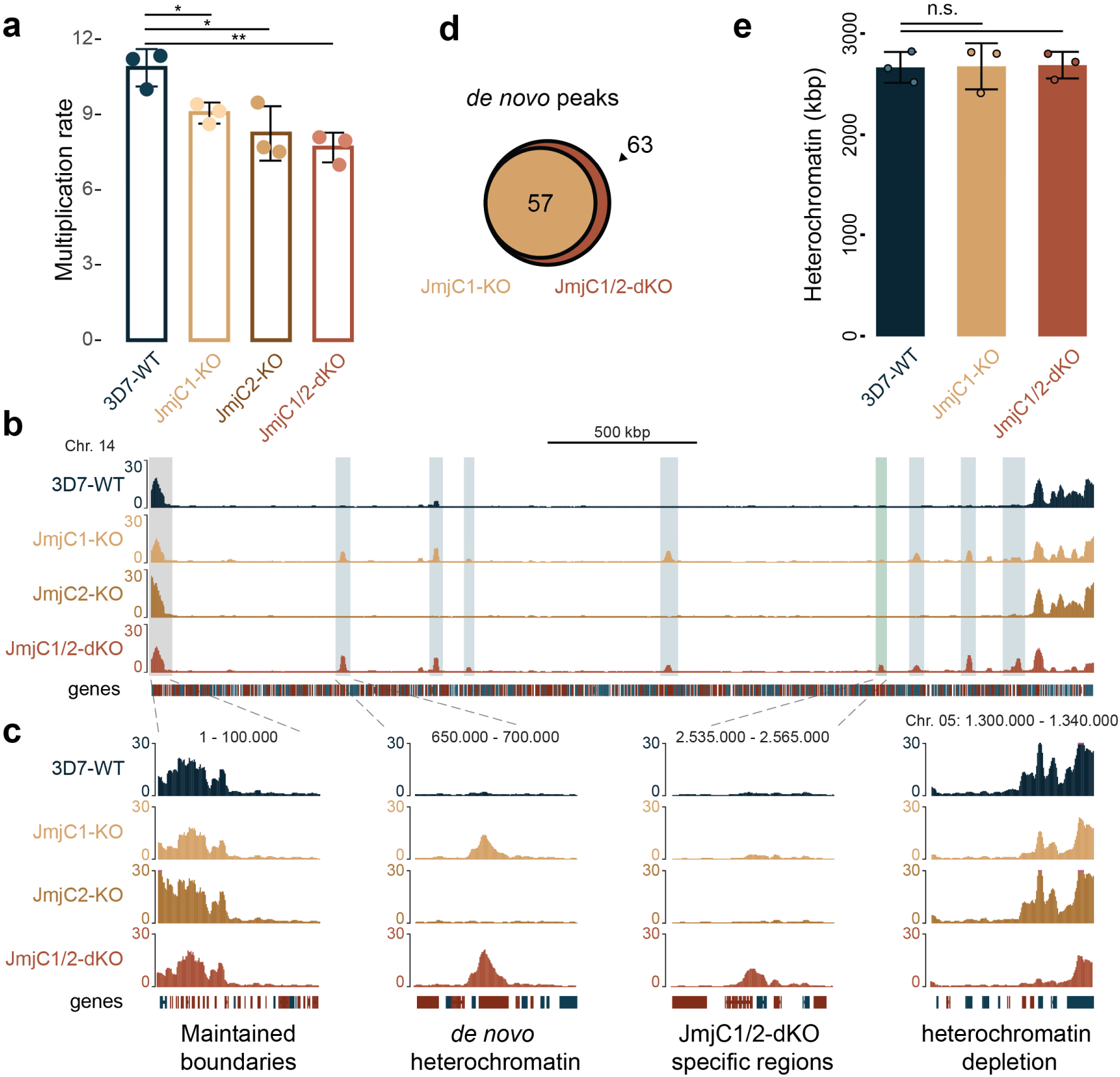
Knockout of PfJmjC1 leads to heterochromatin rearrangement. **a)** Multiplication rate of 3D7 wild type (WT), 3D7/JmjC1-KO, 3D7/JmjC2-KO and 3D7/JmjC1/2-dKO parasites. Values represent the mean of three independent experiments, with dots representing the individual values and error bars defining the standard deviation. Statistical significance was accessed using unpaired two-sided t-tests with Welch’s correction. P-value is shown as * < 0.05 and ** < 0.01. **b)** PfHP1 CUT&Tag read occupancy tracks on chromosome 14 (average of three replicates). Areas of increased heterochromatin occupancy in the mutants compared to 3D7 wild type (WT) are highlighted with a blue hue, those specific to the PfJmjC1/2 double KO line are highlighted with a green hue. The area at the distal end depicts a region with reduced CUT&Tag signal intensity, which for this particular region is further exuberated by the deletion of the chromosome end in a subset of the parasites in the double mutant (Extended Data Fig. 5e). **c)** Left-to-right: Zoom-ins of maintained heterochromatin boundaries, de novo heterochromatin formations in both the JmjC1-KO and JmjC1/2-dKO mutants, de novo formations specific to the JmjC1/2-dKO mutant and areas of heterochromatin depletion to varying degrees in both KO mutants. **d)** Venn diagram depicting the overlap between de novo heterochromatin regions in PfJmjC1 single and PfJmjC1/2 double KO mutants. **e)** Combined length of all heterochromatic regions called in the different strains. Values represent the mean of three independent replicate experiments, with dots representing the individual values and error bars defining the standard deviation. Statistical significance was accessed using unpaired t-test with Welch’s correction (n.s. = not significant).

In order to test whether PfJmjC1 and/or PfJmjC2 are involved in shaping the heterochromatin landscape, we profiled heterochromatic regions in both wild type and the three KO parasite lines utilizing our recently established CUT&Tag protocol^45^. We did not observe heterochromatin spreading from existing domains in either the 3D7/JmjC1-KO, 3D7/JmjC2-KO or 3D7/JmjC1/2-dKO mutant parasites (Fig. 2b, grey hue, Extended Data Fig. 4b). However, we noticed extensive formation of small heterochromatic domains within euchromatic regions in 3D7/JmjC1-KO and 3D7/JmjC1/2-dKO parasites (Fig. 2b-c, blue hue; from now on referred to as *de novo* regions), while 3D7/JmjC2-KO showed no major differences in heterochromatin occupancy compared to the wild type control (Fig. 2c, Extended Data Fig. 4e). In the 3D7/JmjC1/2-dKO mutant, we identified 63 *de novo* heterochromatin regions that are absent in 3D7 wild type parasites, of which 57 were also found in the 3D7/JmjC1-KO mutant and six were only observed in the double mutant (Fig. 2d, Extended Data Fig. 4a, Extended Data Table 1). In parallel to the formation of extra heterochromatin domains, heterochromatin occupancy was reduced at some chromosome ends in both 3D7/JmjC1-KO and 3D7/JmjC1/2-dKO parasites (Fig. 2c, Extended Data Fig. 4d). In fact, the combined length of heterochromatic regions in wildtype and 3D7/JmjC1-KO and 3D7/JmjC1/2-dKO parasite strains was largely comparable to wild type (Fig. 2e). Together, these data show that PfJmjC1, but not PfJmjC2, is centrally involved in heterochromatin organization and preventing heterochromatin formation over euchromatic genes in asexual blood stage parasites. Furthermore, they suggest that loss of PfJmjC1 function leads to rearrangements in the heterochromatin landscape rather than an absolute increase in the amount of heterochromatin in the parasite epigenome.

### Knockout of PfJmjC1 leads to rearrangement of the 3D nuclear architecture

While heterochromatin spread has been shown to occur linearly along the chromosome in other organisms ^46–48^ and *P. falciparum* ^8,49^, it is possible that heterochromatin could also spread to distant regions of the genome via long-range DNA-DNA contacts. To determine if the *de novo* heterochromatin regions found in the 3D7/JmjC1/2-dKO parasites form contacts with existing heterochromatic regions, we performed a high-resolution chromosome conformation capture experiment (Micro-C ^50^) on 3D7 wild type and 3D7/JmjC1/2-dKO parasites (displaying the most prominent heterochromatin rearrangement). We found that the *de novo* heterochromatin regions identified in the mutant do not interact with the subtelomeric or intrachromosomal heterochromatin domains in wild type parasites, suggesting that formation of heterochromatin over these new sites is not due to heterochromatin spreading via preexisting long-range DNA-DNA contacts (Fig. 3a,c; Extended Data Figure 5a,b). However, in the 3D7/JmjC1/2-dKO parasites, a subset of *de novo* heterochromatinised regions showed increased contact frequency with existing heterochromatin regions (Fig. 3c; Extended Data Fig. 5a,b,d; Extended Data Table 2). For example, two *dynein* loci on chromosome 7 (PF3D7_0718000, PF3D7_0729900) that gained PfHP1 occupancy in the mutants show new intra-chromosomal contacts with subtelomeric and central virulence gene-containing PfHP1-demarcated domains on chromosome 7, seemingly forming a new heterochromatin compartment in the nucleus (Fig. 3a). Conversely, some regions of heterochromatin loss, such as the distal end of chromosome 5 (Fig. 3b), show a loss of contacts with other heterochromatic regions in 3D7/JmjC1/2-dKO parasites (Fig. 3b,d). In addition, intra-chromosomal heterochromatin – heterochromatin interactions appear to be generally more frequent in the 3D7/JmjC1/2-dKO line compared to 3D7 wild type parasites (Fig. 3a, Extended Data Figure 5a, Extended Data Table 2).

**Fig. 3:**
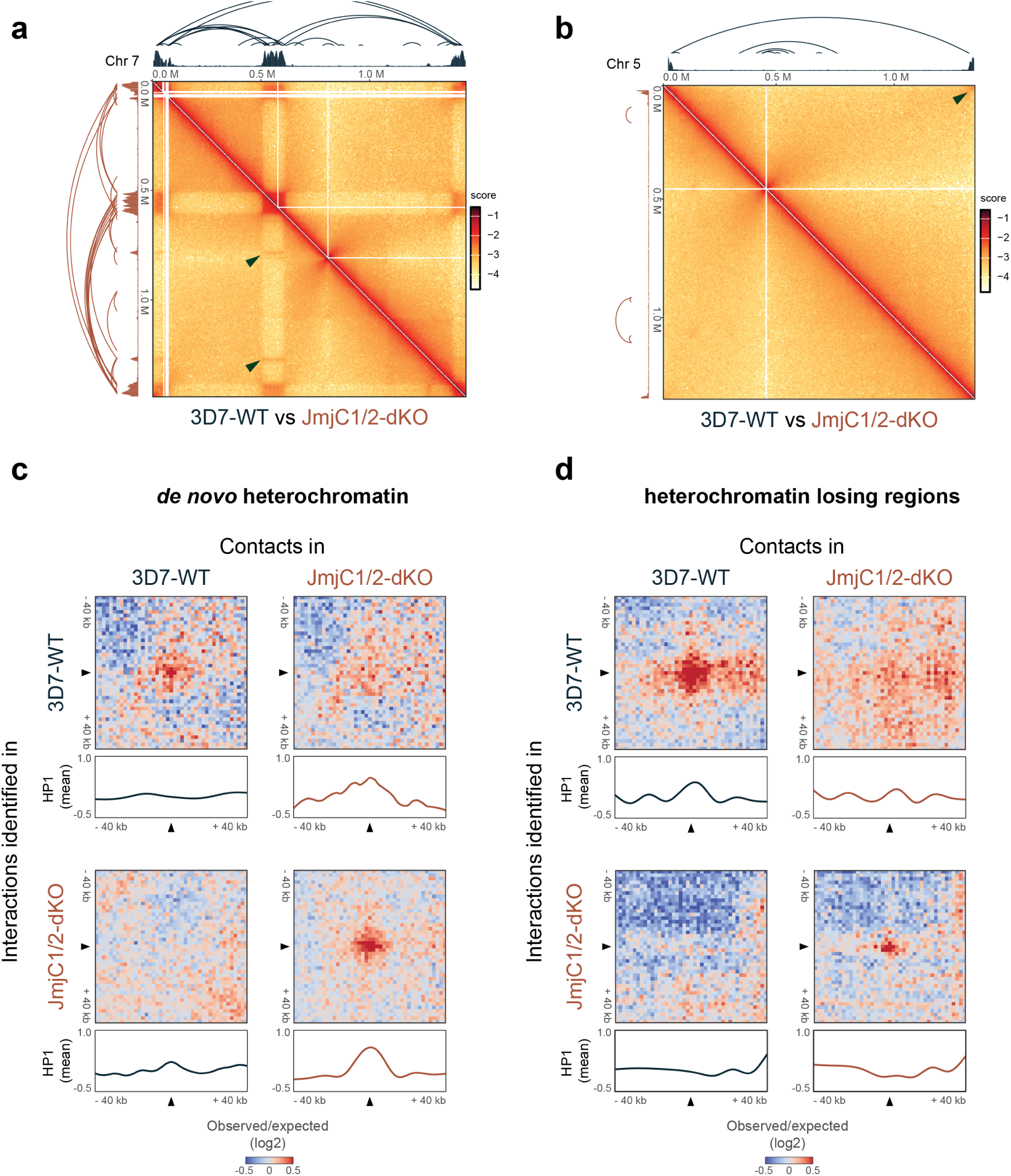
Knockout of PfJmjC1 leads to re-organization of the 3D nuclear architecture. **a,b)** Micro-C contact map of chromosome 7 (a) and chromosome 5 (b) in 3D7 wild type (top right) and PfJmjC1/2-dKO (bottom left) parasites (5kb resolution, normalized interaction frequency) generated from four separate replicates. Color scales are shown on the right. PfHP1 CUT&Tag tracks and identified interactions for 3D7 wild type (dark blue) and 3D7/JmjC1/2-dKO (red) parasites are displayed on the left side and on top, respectively. Strain-specific interactions in the 3D7/JmjC1/2-dKO and 3D7 wild type (WT) parasites are marked with green arrowheads. **c,d)** Off-diagonal Micro-C contact maps, aggregated over long-range interactions (±40kb, 1kb resolution, Extended Data Table 2) in 3D7 wild type (WT) (left) or 3D7/JmjC1/2-dKO (right) parasites. The top row shows long-range interactions identified in 3D7 wild type (WT) parasites with at least one anchor in de novo heterochromatinised (c) or heterochromatin losing regions (d) in the 3D7/JmjC1/2-dKO parasite strain. The bottom row shows long-range interactions identified in 3D7/JmjC1/2-dKO with at least one anchor in de novo heterochromatin (c) or heterochromatin losing regions (d) in the 3D7/JmjC1/2-dKO parasite strain. Color scales indicate log2(observed/expected interaction frequency). Aggregated HP1 CUT&Tag (see Fig. 2b) signals from 3D7 wild type (WT) and 3D7/JmjC1/2-dKO parasites are shown below the Micro-C maps.

In conclusion, Micro-C analysis of the 3D7/JmjC1/2-dKO mutant highlight substantial changes in the nuclear architecture as a consequence of heterochromatin rearrangement.

### *De novo* heterochromatin formation in PfJmjC1 knockout mutants mainly occurs over GC-rich non-expressed and/or non-essential genes

To determine why particular regions of the genome are targeted for *de novo* heterochromatin formation, we characterized these regions in 3D7/JmjC1/2-dKO parasites, which are 5-10 kb long with a median width of 7 kb (Fig. 4a). Notably, we rarely observed regions shorter than 5 kb, suggesting that there might be a minimum size for a newly formed heterochromatin domain to be stable. These regions do not appear to be enriched for any particular DNA motif(s) that could explain specific recruitment of heterochromatin formation to these regions via, for example, DNA-binding proteins (Extended Data 5c). However, the fact that we did not detect significantly enriched motifs could be due to the limited number of *de novo* heterochromatin regions (n=63) and hence the limited power of finding significantly enriched motifs using the HOMER algorithm. Interestingly, however, *de novo* heterochromatinised regions have a significantly higher GC content (median 21%) than randomly selected euchromatic regions (median 18,6%) and comparable to some regions heterochromatic in wild type parasites (median 21,7%, Fig. 4b).

**Fig. 4:**
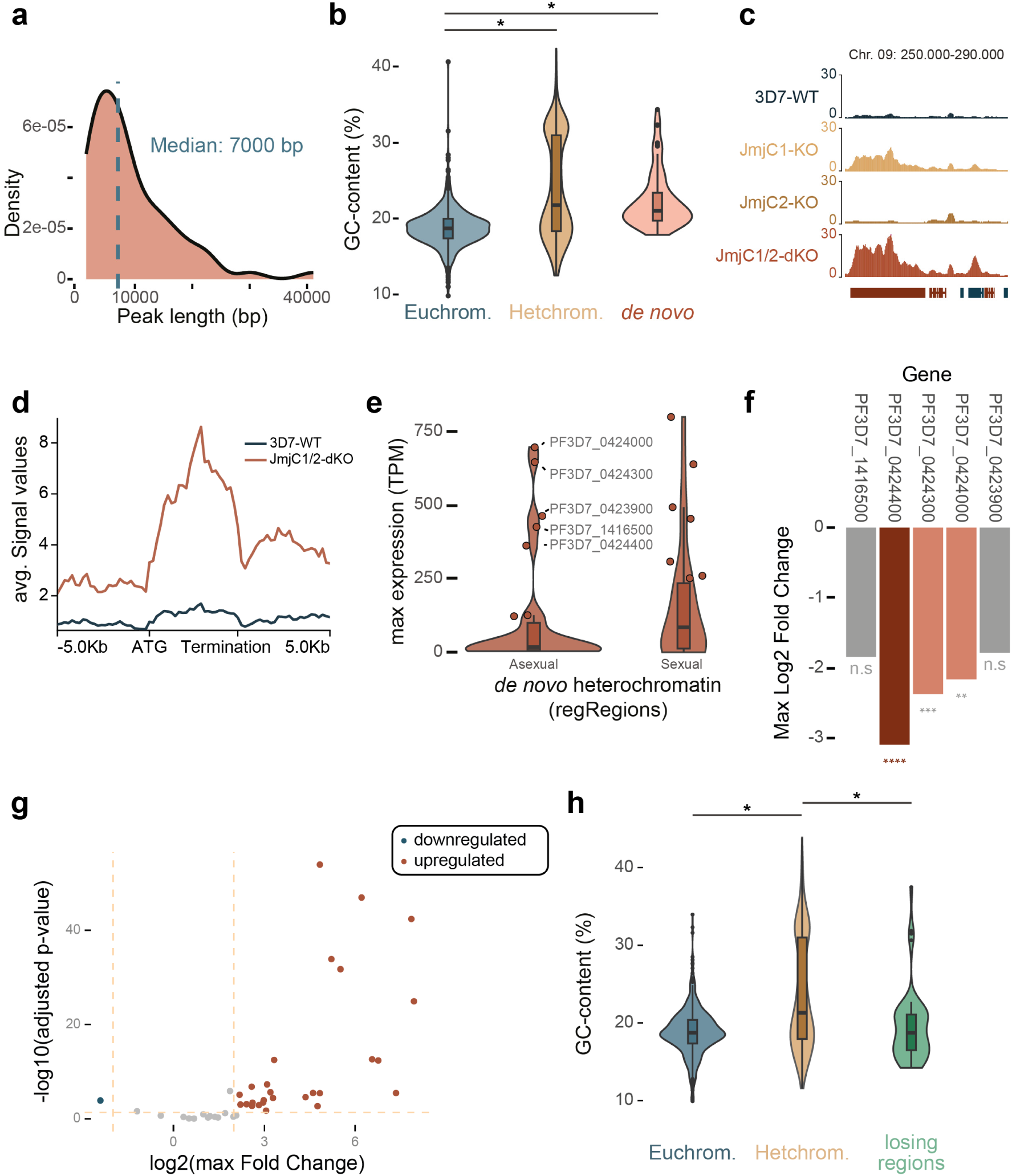
De novo heterochromatin formation in PfJmjC1 mutants occurs on GC-rich gene bodies and lowly expressed genes. **a)** Peak length distribution of all de novo heterochromatin peaks identified in the PfJmjC1/2 double KO mutant, calculated in 500 bp resolution. Y-axis depicting frequency of peak length. **b)** GC-content in de novo heterochromatin regions of the PfJmjC1/2 double KO mutant compared to that of size-matched euchromatic and heterochromatic control regions. Differences between euchromatic control and de novo-formed heterochromatic regions and between euchromatic and heterochromatic control regions are significant as determined by Wilcoxon signed-rank test (p-value < 2.2e-16 for both comparisons). Boxplots show quartiles 1 to 3, median is indicated as a thick line. **c)** Zoom in of representative PfHP1-CUT&Tag read occupancy track to a de novo region on chromosome 9. Average of three replicates. **d)** Average heterochromatin occupancy profile of all 77 genes gaining heterochromatin on their gene bodies in the PfJmjC1/2 double KO compared to 3D7 wild type (WT) parasites. **e)** Maximal expression in asexual replication defined in tag per million reads (TPM) of 30 genes with increased heterochromatin signal in their regulatory regions (-1000 bp ATG +500), excluding PF3D7_0905400 which had an expression value of 5736,93. Values are based on RNA-seq data from Toenhake et. al^51^. Boxplots show quartiles 1 to 3, median is indicated as a thick line. **f)** Max log2 fold change in abundance of the transcripts highlighted in panel **e**, assessed by DESeq2 between 3D7 wild type (WT) and 3D7/JmjC1/2-dKO parasites. Red-tinted genes are significantly downregulated. Adjusted p-value is shown as ** < 0.01 and *** < 0.001. **g)** GC-content of heterochromatin-depleted regions in the PfJmjC1/2 double KO mutant compared to that of size-matched euchromatic and heterochromatic control regions. Significant differences were determined by Wilcoxon signed-rank test (p-value < 2.2e-16 for both comparisons). Boxplots show quartiles 1 to 3, median is indicated as a thick line. **h)** Volcano plot of maximum fold change over measured time points and corresponding adjusted p-value as determined by DESeq2 for all genes losing heterochromatin over their regulatory regions. Upregulated genes (red dots) were defined as log2(foldchange) > 2 and adjusted p-value < 0.05, downregulated genes (blue dots) as log2(foldchange) < -2 and adjusted p-value < 0.05.

**Fig. 5:**
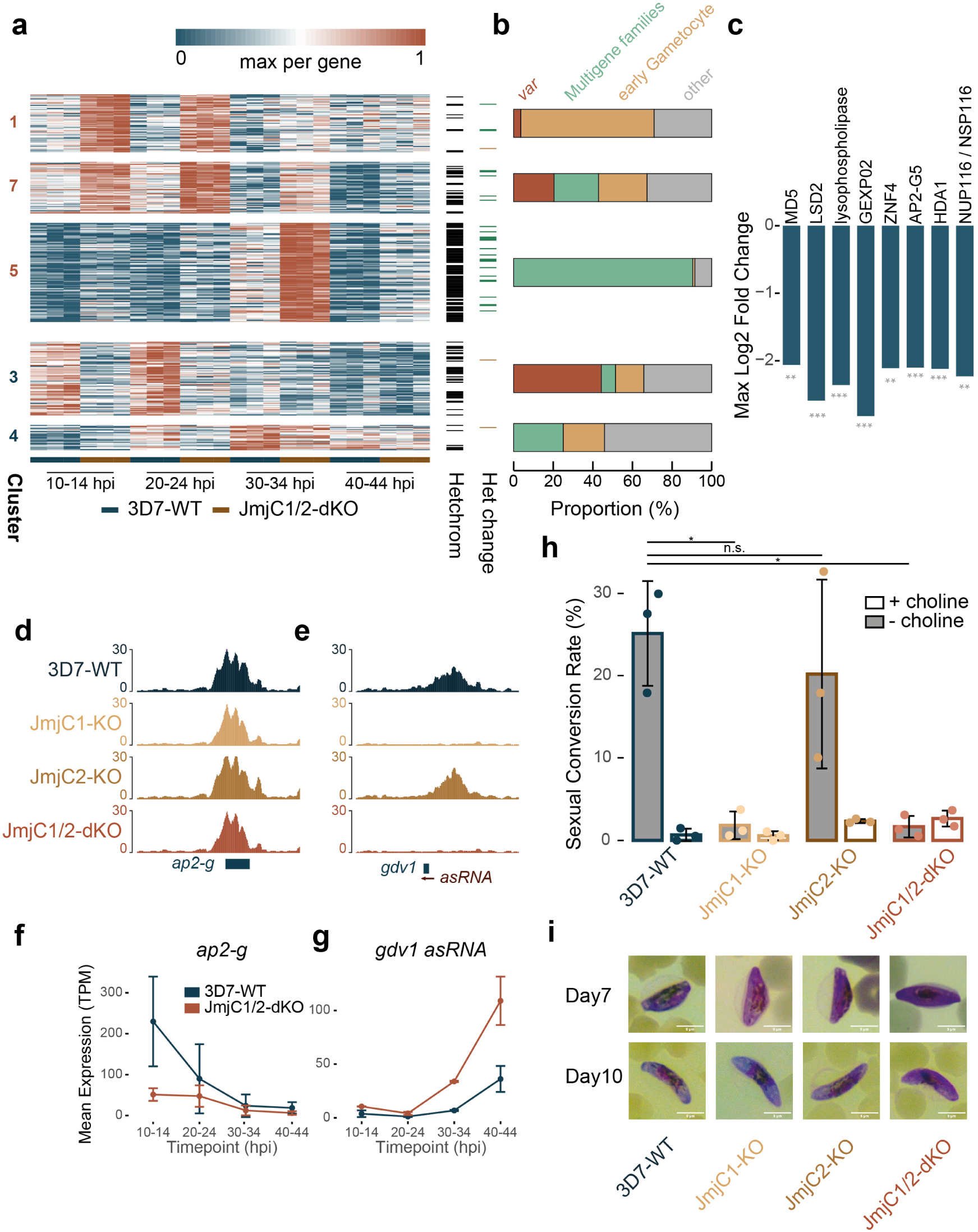
PfJmjC1 is required for proper regulation of gene expression and induction of sexual commitment in response to nutrient depletion. **a)** Heatmap of RNA-seq data of genes with altered transcript abundance between 3D7 wild type and 3D7/JmjC1/2-dKO parasites. Transcript abundance values represent three replicate experiments per condition/timepoint and have been scaled to their maximum expression value per gene. Clustering was performed with k-means and heterochromatin state in wild type parasites is indicated by a black bar (Fraschka et. al, 2018)^8^. Changes in heterochromatin occupancy comparing wild type to 3D7/JmjC1/2-dKO parasites are shown as green (heterochromatin-depleted) or brown (heterochromatin-gaining) bars. **b)** Proportions of genes in each cluster that belong to var (red), other multigene families (green), early gametocyte genes (yellow) or other genes (grey). **c)** Highest log2 fold change of all timepoints of select early gametocyte genes, calculated on three independent replicate experiments by Deseq2. Adjusted p-value is shown as ** < 0.01 and *** < 0.001. **d,e)** PfHP1 CUT&Tag read occupancy tracks over the pfap2-g (d) and **e)** the gdv1 (e) locus. **f,g)** Mean transcript abundance of ap2-g(f) and gdv1 asRNA (g) in TPM . Error bars define the standard deviation. **h)** Sexual conversion rates of 3D7 wild type, 3D7/JmjC1-KO, 3D7/JmjC2-KO and 3D7/JmjC1/2-dKO parasites cultured in minimal fatty acid medium supplemented with 2 mM choline chloride (+choline) (baseline sexual commitment rates) (open bars) or not (-choline) (sexual commitment-inducing conditions) (filled bars) as assessed by anti-Pfs16 IFAs. Values represent the mean of three independent experiments. Dots represent individual values and error bars define the standard deviation. Significance was calculated using a one-way Anova t-test. p-value is shown as * < 0.05. i) Representative images of Hemacolour-stained blood smears of of 3D7-WT, 3D7/JmjC1-KO, 3D7/JmjC2-KO and 3D7/JmjC1/2-dKO gametocytes.

We found that *de novo* heterochromatic domains usually overlap with one to three genes and are mostly deposited on gene bodies rather than upstream regulatory regions (77 genes have enriched heterochromatin on gene bodies, 31 genes at regulatory regions) (Fig. 4c, Extended Data Table 1). Accordingly, the distribution of heterochromatin signal over all heterochromatin-gaining gene bodies in 3D7/JmjC1/2-dKO parasites shows a slightly higher heterochromatin occupancy towards the middle and 3’ end of affected genes (Fig. 4d). Notably, most genes that have increased heterochromatin signal within their regulatory region (-1000 bp to +500 bp surrounding the translation start site) are lowly expressed in 3D7 wild type asexual parasites (Fig. 4e, left violin plot) and tend to be non-essential for asexual replication (Extended Data Fig. 5d). Furthermore, most affected genes are expressed at and relevant for later life cycle stages such as gametocytogenesis and ookinete formation (Fig. 4e, Extended Data Table 1).

To assess potential repressive properties of *de novo* heterochromatin on gene transcription, we performed RNA-seq experiments on synchronous 3D7 wild type and 3D7/JmjC1/2-dKO parasites at four different time points during intra-erythrocytic parasite development [10-14, 20-24, 30-34 and 40-44 hours post invasion (hpi)] (Extended Data Fig. 6a). Out of the five heterochromatin-gaining genes that are highly expressed in asexual parasites (transcripts per million (TPM) > 200), all showed downregulation in 3D7/JmjC1/2-dKO parasites at their respective peak expression time points, three of which at statistically significant levels (PF3D7_0424000, PF3D7_0424300, PF3D7_0424400) (Fig. 4f). Interestingly, all of these genes were dispensable in asexual parasites^38^ (Extended Data Table 1).

In contrast to heterochromatin-gaining genes, genes located in regions with reduced heterochromatin levels in the PfJmjC1 mutants tend to encode exported proteins including members of the PfEMP1, PHISTs and RIF protein families and were upregulated in 3D7/JmjC1/2-dKO compared to 3D7 wild type parasites (Extended Data Table 3). Specifically, 27 out of 42 heterochromatin-depleted genes were upregulated in the 3D7/JmjC1/2-dKO mutant (Fig. 4g). Interestingly, heterochromatin-depleted genes are typically located in heterochromatic regions with a lower GC content (median 18,7%) compared to size-matched random heterochromatic regions (median 21,3%), which rather mirrors the average GC content of euchromatic regions (median 18,7 %) (Fig. 4h).

Overall, these analyses highlight that *de novo* heterochromatin formation in the 3D7/JmjC1-KO and 3D7/JmjC1/2-dKO parasites is unlikely to be a targeted process, i.e. not guided by DNA-binding proteins but can occur over non-expressed and/or non-essential euchromatic genes. Furthermore, our data is consistent with the notion that *de novo* heterochromatin formation preferentially occurs over several kilobase long GC-rich euchromatic regions, while loss of heterochromatin primarily occurs in heterochromatic regions with similar GC content.

### PfJmjC1 knockout affects expression of heterochromatic genes and decreases sexual conversion

Transcriptome-wide analysis of the RNA-seq data revealed 325 and 112 significantly up- and downregulated genes, respectively, at any of the four timepoints assessed (fold-changes of >2 or < -2; adjusted p-values < 0.01) (Extended Data Table 3). Differentially expressed genes were clustered into seven groups by k-means clustering. Clusters 1, 5 and 7 contain genes that display higher transcript abundance in ring (10/20 hpi) or trophozoite (30 hpi) stages of 3D7/JmjC1/2-dKO mutant parasite. Clusters 3 and 4 contain genes with lower transcript abundance in rings and trophozoite stages of the mutant (Fig. 4a). The genes in clusters 2 and 6 show somewhat less clear upregulation pattern at various stages (Extended Data Fig. 6b).

Interestingly, genes that are heterochromatic in wild type parasites were overrepresented among the deregulated genes (∼5-fold, hypergeometric test: p-value = 6.1e-81) and, particularly in cluster 5, consist of genes predominantly from the *rifin*, *stevor* and *pfmc-2tm* gene families (Fig. 4a,b, Extended Data Table 3). Consistent with the upregulation of these genes, clusters 1, 5 and 7 further contained almost all genes with reduced heterochromatin occupancy in the 3D7/JmjC1/2-dKO mutant. Yet not all genes with higher transcript abundance display a significant change in heterochromatin occupancy; deregulation of these genes could hence be the consequence of smaller changes in heterochromatin occupancy undetectable in a bulk analysis. Notably, *var* genes (encoding the major cell surface antigen PfEMP1) were prominently present amongst both up- and down-regulated genes (cluster 7 and cluster 3), showing that depletion of PfJmjC1 and PfJmjC2 does not lead to global upregulation of *var* genes (Extended Data Fig. 6c) in line with a previous report^20^. Changes in the expression of heterochromatic multigene families between mutant and wild type parasites could in part be explained by clonally variant gene expression^2^ and is most likely the case for the *var* genes. On the other hand, *rifin*, *stevor* and *pfmc-2tm* genes are more often upregulated than downregulated in mutant parasites and hence their deregulation is more likely caused by PfJmjC1/2 depletion and heterochromatin rearrangements.

Interestingly, next to the deregulation of members of heterochromatic multigene families, we also observed a altered transcript abundance of 95 gametocytogenesis-related genes potentially indicating altered gametocyte numbers and/or commitment dynamics between wild type and 3D7/JmjC1/2-dKO parasites (Extended Data Table 3). In particular, many of the immediate early markers of gametocytogenesis, including *pfap2-g*^52,53^*, pfap2-g5*^54^*, pfgexp02*^55,56^ and *pfnup116*^57^, displayed lower abundance in the 3D7/JmjC1/2-dKO mutant parasites (Fig. 4c). We therefore closely examined heterochromatin distribution over the *pfap2-g* locus but did not observe any marked changes in 3D7/JmjC1/2-dKO compared to wild type parasites (Fig. 4d,f). Most notably, however, we observed a complete loss of heterochromatin downstream of the *gdv1* gene along with upregulation of the *gdv1* long non-coding antisense RNA (*gdv1* asRNA)^58^ in the 3D7/JmjC1/2-dKO mutant (Fig. 4e,g). Since fine-tuned expression of the *gdv1* asRNA has been shown to be relevant for regulating GDV1 expression and in turn commitment to gametocytogenesis^14,15^, we assessed sexual conversion rates in the PfJmjC mutant parasites.

All three PfJmjC KO lines were able to produce gametocytes at the typically low baseline rates obtained under standard *in vitro* culture conditions (Fig. 4h, Extended Data Fig. 7a). Furthermore, gametocyte development seemed to progress normally as judged by visual inspection and female:male sex ratios were comparable in all mutants and 3D7 wild type control (Fig. 4i, Extended Data Fig. 7b). Strikingly, however, only 3D7 wild type and 3D7/JmjC2-KO parasites show the expected increase in sexual conversion rates upon exposure to lysophosphatidylcholine-/choline-free minimal fatty acid medium^59^ while this capacity is completely abolished in the 3D7/JmjC1-KO and 3D7/JmjC1/2-dKO parasites (Fig. 4h).

Collectively, our comprehensive analysis shows that PfJmjC1/2-dKO parasites have an altered gene expression profile in concordance with a changed heterochromatin landscape, mainly affecting heterochromatic multigene families. Furthermore, we demonstrate that 3D7/JmjC1-KO and 3D7/JmjC1/2-dKO parasites are unable to induce gametocytogenesis in response to choline depletion, while basal sexual conversion rates are similar between wildtype and mutant parasites.

## Discussion

In this study, we demonstrate that the histone demethylase PfJmjC1 is essential for proper heterochromatin distribution, especially in preventing heterochromatin formation in euchromatic regions. PfJmjC1 interacts with PfHP1, the main constituent of heterochromatin. In 3D7/JmjC1-KO and 3D7/JmjC1/2-dKO parasites, heterochromatin is redistributed, leading to frequent *de novo* heterochromatin formation as well as heterochromatin loss at some chromosome ends. Heterochromatin gain does not occur at loci that form preexisting (i.e. prior to deletion of PfJmjC1) long-distance interactions with other heterochromatic regions, but eventually results in rearrangement of nuclear architecture, leading to increased interactions between *de novo*-formed PfHP1 islands with established heterochromatic regions. Loss of PfJmjC1 leads to decreased gametocyte production rates under nutrient depletion a most probably due to the depletion of heterochromatin over the promoter of the *gdv1* antisense transcript. All in all, we show how PfJmjC1 is imperative to maintain proper heterochromatin distribution and related transcriptional repression as well as life cycle stage transition.

Next to PfHP1, PfJmjC1 interacts with several other proteins (Fig. 1e,f) including ApiAP2 factors and posttranslational modifiers such as the phosphatase PfPPM4, the most highly enriched interaction partner of PfJmjC1. This interaction suggests that their shared targets in heterochromatin may require both demethylation and dephosphorylation for proper cellular function or as a specific posttranslational switch to activate / deactivate certain properties. Alternatively, it could indicate that PfJmjC1 is a target of PfPPM4. Phosphorylation indeed plays an important role in the regulation of HDMs in other organisms (e.g humans)^60^, where it for instance affects the subcellular localization of KDM5A^61^, protein half-life of KDM4C^62^ and complex dissociation in LSD1+8a^63^. Cheng et al. have also shown that the human JmjC domain-containing demethylase KDM3A is phosphorylated upon heat-shock, forming a complex with the transcription factor STAT1 and exhibiting a different chromatin-binding specificity^64^. This opens the possibility of PfJmjC1 specifically binding to certain areas of the genome depending on its phosphorylation state as well as interaction with different ApiAP2 transcription factors. Furthermore, the activity of PfJmjC1 might be altered by certain environmental stimuli via phosphorylation/dephosphorylation and lead to altered heterochromatin organization. This in turn might influence expression of cell surface antigens and/or gametocyte commitment rates, ultimately influencing parasite survival and fine-tuning transmission.

JmjC1 is conserved in its domain structure across several malaria parasite species infecting humans, such as *P. falciparum*, *P. vivax*, *P. malariae*, and *P. knowlesi* (Fig. 1), but no orthologue is present in rodent malaria parasite species like *P. berghei* (www.plasmoDB.org). In contrast, orthologs of both PfJmjC2 and PfJmjC3 are present in *P. berghei*. Intriguingly, the lack of clear PfJmjC1 orthologs in *Plasmodium spp.* that infect rodents coincides with the scarcity of intrachromosomal heterochromatic islands in these species, while in species which have a PfJmjC1 ortholog, intrachromosomal islands are more frequent^8^. Therefore, JmjC1 may have emerged or been retained in malaria parasites infecting apes and humans to prevent the formation of undesired intrachromosomal heterochromatin islands.

*De novo* heterochromatic islands are the most prominent feature in parasites lacking PfJmjC1 and are not present in 3D7/JmjC2-KO parasites. Yet, despite the lack of PfJmjC1 and PfJmjC2 activity, these islands remain rather delineated, and we did not observe further heterochromatin spread neither from these *de novo* nor from existing heterochromatic domains (Fig. 2). This might be due to the essentiality of genes neighboring heterochromatic regions, and hence spreading of heterochromatin to these areas might be lethal for the parasite. This possibility is consistent with the slight growth defect of 3D7/JmjC1-KO and 3D7/JmjC1/2-dKO mutant parasites; parasites in which heterochromatin spread over essential genes occurs die, whereas parasites “keeping” heterochromatin regions confined survive. Alternatively, further spread of heterochromatin may require removal of a second mechanism safeguarding heterochromatin distribution. In fission yeast, for example, only the combined knock-out of the JmjC domain-containing demethylase Epe1 and the histone acetyltransferase Mst2 leads to substantial heterochromatin spreading events^23^. It is thus possible that in the absence of PfJmjC1 , histone acetylation-based activities may prevent further H3K9me3 deposition and local heterochromatin spread. Finally, next to the lack of local heterochromatin spread, it is notable that in the 3D7/JmjC1-KO and 3D7/JmjC1/2-dKO mutants the total amount of heterochromatin remains constant (Fig. 2f), suggesting that heterochromatin gain in euchromatic regions is accompanied by heterochromatin loss in exiting heterochromatic domains. This finding suggests the presence of a limiting factor that prevents excessive spread of heterochromatin. For example, in Epe1 - Mst2 double mutant *S. pombe*, heterochromatin is deposited at the *hp1/swi6* locus, limiting the amount of HP1/SWI6 available for heterochromatin formation^23^. While we did not observe accumulation of heterochromatin at the *pfhp1* locus in 3D7/JmjC1-KO and 3D7/JmjC1/2-dKO parasites (Extended Data Fig. 2e), other self-limiting mechanisms might exist that keep total heterochromatin levels in control.

Formation of *de novo* heterochromatin island lead to substantial rearrangement of the 3D nuclear architecture, providing additional evidence for the link between heterochromatin and long range DNA-DNA interactions. Yet, we did not detect new interaction for all *de novo* heterochromatic regions. While this could be due to the incomplete sampling of interactions by MicroC it could also indicate that a small heterochromatin island alone is not always sufficient to mediate these interactions and there may be other protein factors working with HP1 to facilitate stable 3D contacts.

Transcriptomic changes in the 3D7/JmjC1/2-dKO parasites can to a large degree be explained by the observed heterochromatin rearrangements, as many deregulated genes belong to heterochromatic multigene families (Fig. 4), some of which prominently lose heterochromatin in the mutants (Fig. 3). Generally, many genes encoding exported proteins were upregulated, among which are nine out of the eleven *pfmc-2tm* genes, 65 *rif* and 13 *stevor* genes (Extended Data Table 3). Given the massive deregulation of heterochromatic gene families encoding exported proteins, it is conceivable that the antigen repertoire and/or presentation on the surface of the infected RBC is substantially altered in parasites lacking expression of PfJmjC1. Such changes are not expected to lead to any adverse consequences under *in vitro* culture conditions, consistent with the rather weak growth defect of the PfJmjC mutants, but may be detrimental in natural infections due to increased adaptive-immunity-dependent parasite elimination.

Next to altered expression of cell surface antigens, genes involved in gametocytogenesis are also display differential abundance in the 3D7/JmjC1/2-dKO parasite populations, which caused us to examine gametocyte development. Strikingly, PfJmjC1 mutant parasites have a severely reduced sexual conversion rate under nutrient-depleted, gametocytogenesis-inducing conditions, which is most likely the consequence of the loss of heterochromatin downstream of the *gdv1* locus. According to our current understanding, this heterochromatic island in wild type parasites suppresses the expression of the long non-coding *gdv1* asRNA, which in turn is suppresses expression of GDV1 and activation of PfAP2-G. Variation in the level of heterochromatin over the promoter of the *gdv1* asRNA transcript in various parasite strains has previously been reported to correlate with the amount of gametocytes produced^14,15,58^. Hence, it is tempting to speculate that PfJmjC1 might be involved in regulating the amount of heterochromatin over this locus and indirectly influencing the transmission potential of the parasite under nutrient depletion. Nonetheless, the loss of heterochromatin over this particular region in 3D7/JmjC1-KO and 3D7/JmjC1/2-dKO mutants is intriguing, given that over the rest of the genome multiple new intrachromosomal PfHP1 islands are formed. Hence, it is not unthinkable that upon loss of PfJmjC1, heterochromatin levels increase over the *gdv1* asRNA locus leading to sexual commitment and consequently loss of these parasites from the proliferating population, and the ultimate loss of heterochromatin over this locus in our PfJmjC1 KO mutants might be the consequence of a selection for parasites that do not assemble this heterochromatin island anymore. JmjC1 or its downstream targets could further be involved in the lysophosphatidylcholine-sensing pathway responsible for gametocytogenesis induction under nutrient-depleted conditions. Finally, while gametocyte development in the 3D7/JmjC1-KO and 3D7/JmjC1/2-dKO lines seemed to be unaffected based on gametocyte morphology, given the deregulation of and heterochromatin formation over gametocytogenesis-related genes PfJmjC1 mutant gametocytes might not support transmission to a new host.

In this study, we present the first comprehensive characterization of a JmjC-domain-containing demethylase in the context of heterochromatin organization in malaria parasites. The *de novo* formation of numerous intrachromosomal heterochromatin islands upon deletion of PfJmjC1 is a unique phenomenon that, to the best of our knowledge, has not been reported in any organism. This not only highlights the critical function of PfJmjC1 in preventing undesired heterochromatin formation in euchromatic regions in *P. falciparum*, but also reveals mechanistic principles of heterochromatin formation and will therefore aid further exploration of this critical regulatory process in general. Similarly, the effect of PfJmjC1 depletion on sexual conversion is intriguing and suggests that PfJmjC1 may play a crucial role in the regulatory pathway that triggers sexual commitment in response to environmental stimuli. Whether this effect is solely due to the alteration of the heterochromatic island downstream of *gdv1* or also via alternative processes and whether lack of PfJmjC1 influences parasite transmission beyond sexual commitment, remains to be determined.

## Data and code availability

All raw and processed sequencing data have been submitted to Gene Expression Omnibus (GEO) under the reference numbers GSE306368, GSE306369 and GSE309221.

Code used for analysis of heterochromatin regions, RNA-seq experiments is available at our github page https://github.com/bartfai-lab/JmjCs_heterochromatin-calling_RNAseq_P.falciparum

Proteomic data has been submitted to PRIDE under the accession number PXD066534.

## Acknowledgments

J.G., B.G., T.V. and R.B. have received funding from the EU’s Horizon 2020 research and innovation program (Cell2Cell ITN) under the Marie Skłodowska-Curie grant agreement number 860875. This work was further supported by Swiss National Science Foundation grants BSCGI0_157729 and 310030_220001. Work in the Bartfai lab has been supported by a ERC Synergy grant (# 101118536). We want to thank Sebastian Baumgarten for his valuable input regarding the Micro-C experiments.

## Author contributions

Conceptualization, J.G., T.V. and R.B.; investigation, J.G., M.W., P.S., A.K., B.G., D.K., T.B. and T.R.; methodology, J.G. and P.S.; formal analysis, J.G., M.W., and P.S.; visualization, J.G. and M.W.; data curation, J.G., P.S. and A.K.; writing – original draft, J.G., T.V. and R.B.; writing – review & editing, J.B.; funding acquisition, T.V. and R.B.; supervision, J.B., T.V and R.B.

## Methods Cell culture

*P. falciparum* intraerythrocytic parasites were cultured in human RBCs (Sanguin Nijmegen, Netherlands and Blutspendezentrum Zürich, Switzerland) at 37 °C under low oxygen conditions (3% O_­_, 4% CO_­_and 93% N_­_) at 5% hematocrit in RPMI 1640 medium supplemented with 25 mM HEPES, 0.2% NaHCO_­_, 200 µM hypoxanthine (Merck, H9377) and 10% human serum (Sanguin Nijmegen, Netherlands) or 0.5% Albumax II (Gibco, #11021037) supplemented with 2 mM choline chloride. Parasites were grown either in the absence of antibiotics or in the presence of 0.1g/l neomycin.

In order to obtain synchronized parasites populations, cultures were subjected to repeated sorbitol-based lysis of remodeled RBCs^65^. Specifically, RBCs were pelleted, resuspended in seven volumes of 5% sorbitol and incubated at 37 °C for 10 min. Treated RBCs were washed once with culture medium and cultured further under standard culture conditions.

### Cloning of CRISPR/Cas9 transfection constructs

All transgenic cell lines were generated using our previously published mother plasmids for gene tagging (two-plasmid approach; CRISPR/Cas9 plasmid pH_gC and donor plasmid pD) or gene disruption (all-in-one plasmid approach; CRISPR/Cas9 plasmid p_gC)^15^. To engineer the 3D7/JmjC1-GFP line, we cloned the pH_gC_*jmjc1-gfp* plasmid carrying expression cassettes for the *Streptococcus pyogenes* Cas9 enzyme (SpCas9), human dihydrofolate reductase resistance marker (hd*hfr*) and a small guide RNA (sgRNA) targeting the *jmjc1* coding sequence, and pD_*jmjc1-gfp* containing a sequence assembly designed to tag *pfjmjc1* in frame with *gfp* via homology-directed repair. pH_gC_*jmjc1-gfp* was obtained by inserting annealed complementary oligonucleotides (sgJ1_1F and sgJ1_1R) encoding a sgRNA targeting the *pfjmjc*1 3’ end (gaacacacctacactatatg) into BsaI-digested pH_gC using T4 DNA ligase. pD_*jmjc1-gfp* was generated by a four-fragment Gibson Assembly reaction^66^ joining (1) the pD plasmid backbone amplified from pUC19 using primers PCRA_F and PCRA_R; (2) a 548 bp 5’ HB representing the 3’ end of the *pfjmjc1* coding sequence amplified from a synthetic recodonised *pfjmjc1* sequence (Genscript) using primers J1_5HB_F and J1_5HB_R; (3) the *gfp* sequence amplified from pD_*ap2g-gfp-dd-glms*^15^ using primers gfp_F and gfp_R; and (4) a 3’ HB encompassing the 870 bp directly downstream of the *pfjmjc1* STOP codon amplified from 3D7 gDNA using primers J1_3HB_F and J1_3HB_R.

The 3D7/JmjC1-KO and 3D7/JmjC2-KO cell lines were created using the all-in-one mother plasmid p_gC that encodes expression cassettes for the *Streptococcus pyogenes* Cas9 enzyme (SpCas9) and a sgRNA. To generate the p_gC_*jmjc1-ko_hdhfr* plasmid, annealed complementary oligonucleotides (sgJ1_2F and sgJ1_2R) encoding a *jmjc1*-specific sgRNA (ggtggaacatcatatatggg) were first inserted into BsaI-digested p_gC using T4 DNA ligase to obtain p_gC_*jmjc1*. The final p_gC_*jmjc1-ko_hdhfr* plasmid was then generated via a four-fragment Gibson Assembly reaction joining (1) the BamHI/HindIII-digested p_gC_*jmjc1* plasmid; (2) a 485 bp 5’ HB mapping to the 5’ end of the *pfjmjc1* coding sequence, amplified from 3D7 gDNA using primers J1KO_5HB_F and J1KO_5HB_R; (3) a h*dhfr* expression cassette controlled by the *P. falciparum* calmodulin gene promoter (*cam 5’*) and the *P. berghei dhfr-ts* terminator (*pbdt 3’*), amplified from pH_gC^15^ using primers hdhfr_F and hdhfr_R; and (4) a 588 bp 3’ HB mapping to the 3’ end of the *pfjmjc1* coding sequence, amplified from 3D7 gDNA using primers J1KO_3HB_F and J1KO_3HB_R. To generate the p_gC_*jmjc2-ko_bsd* plasmid, annealed complementary oligonucleotides (sgJ2_F and sgJ2_R) encoding a *pfjmjc2*-specific sgRNA (tataaataaattggtttgtc) were cloned into p_gC as explained above (p_gC_*jmjc2*). The final p_gC_*jmjc2-ko_bsd* plasmid was then generated via a four-fragment Gibson Assembly reaction joining (1) the BamHI/HindIII digested p_gC_*jmjc2* plasmid; (2) a 416 bp 5’ HB mapping to the 5’ end of the *pfjmjc2* coding sequence, amplified from 3D7 gDNA using primers J2KO_5HB_F and J2KO_5HB_R; (3) a blasticidin deaminase (*bsd*) expression cassette controlled by the *cam 5’* and *pbdt 3’* regulatory regions, amplified from p_gC_*map1-ko-bsd*^67^ using primers bsd_F and bsd_R; and (4) a 565 bp 3’ HB mapping to the 3’ end of the *pfjmjc2* coding sequence, amplified from 3D7 gDNA using primers J2KO_3HB_F and J2KO_3HB_R. All oligonucleotides used for plasmid cloning are listed in Supplementary Table S1.

### Parasite transfection

To obtain the 3D7/JmjC1-GFP line, 3D7 wild type parasites were co-transfected with 50 µg each of the pH_gC-*jmjc1-gfp* CRISPR/Cas9 and pD_*jmjc1-gfp* donor plasmid as described^15^. On the day after transfection, the culture medium was exchanged and 4 nM WR99210 (WR) added for six subsequent days, with daily medium replacement. From day seven onwards, WR was removed from the culture medium. Once stably growing transgenic populations were obtained (3-4 weeks after transfection), correct editing of the *pfjmjc1* locus and absence of wild type parasites was confirmed by PCRs on genomic DNA (gDNA , Extended Data Figure 2a,b). All oligonucleotides used for diagnostic PCRs are listed in Supplementary Table S1.

3D7/JmjC1-KO parasites were obtained by transfecting 3D7 wild type parasites with 100 µg of the p_gC_*jmjc1-ko_hdhfr* CRISPR/Cas9 all-in-one plasmid followed by selection of transgenic parasites in the constant presence of 4 nM WR added one day after transfection. 3D7/JmjC2-KO and 3D7/JmjC1/2-dKO parasites were obtained as explained above after transfection of the p_gC_*jmjc2-ko_bsd* plasmid into 3D7 wild type and 3D7/JmjC1-KO parasites, respectively, followed by selection on 2.5 µg/ml BSD-S-HCl. Successful disruption of the *pfjmjc1* and *pfjmjc2* genes in the respective knockout lines were validated by CUT&Tag read occupancy maps, showing no reads mapping to the deleted regions (Extended Data Fig. 3c,d).

### Quantification of parasite multiplication rates

Parasite multiplication rates were determined using flow cytometry. Synchronised parasite cultures were maintained for one replication cycle under standard culture conditions. Parasitemia was measured at the start of the experiment (day 1, 16-24 hpi) and after reinvasion (day 3, 16-24 hpi). Each sample was stained with 1x SYBR-Green I (Invitrogen #S7563), incubated for 20 min in the dark, and washed once in PBS. Using a MACS Quant Analyzer 10, 100,000 events per sample were measured and analyzed using the FlowJo_v10.6.1 software. The proportion of iRBCs was determined based on the SYBR Green I signal intensity. The gating strategy is shown in Extended Data Fig. 7.

### Sexual conversion assays and gametocytogenesis

Synchronized late ring stage cultures (16–24 hpi, 1-1.5% parasitemia) were exposed for 32 hours to minimal fatty acid medium (mFA), containing 0.39% fatty acid-free BSA (Sigma #A6003) instead of 0.5% Albumax II plus 30 μM each of the two essential fatty acids oleic and palmitic acid (Sigma #O1008 and #P0500), supplemented with either 2 mM choline chloride (sexual commitment-suppressing conditions) or 100 µM choline chloride (sexual commitment-inducing conditions) as described^59^. Two methods were used to determine sexual conversion rates: I) Parasites were cultured for another 30 hours before methanol-fixed thin blood smears were prepared at 30–38 hpi and IFAs performed using mouse mAb α-Pfs16^68^ and DAPI. Sexual conversion rates were determined by quantifying the proportion of Pfs16/DAPI double-positive parasites among all DAPI-positive parasites (minimum of 150 parasites scored per sample) (Fig. 5h); II) Parasites were stained for 20 min at 37 °C in the dark with 5 µg/ml Hoechst (Merck #94403) to determine the parasitemia after reinvasion (day 1) using flow cytometry. For the next four days the parasites were cultured in the presence of 50 mM N-acetylglucosamine (GlcNAc) to eliminate asexual parasites^69^. To determine the gametocytemia on day 5, samples were stained for 45 min at 37°C in the dark with 200 nM BioTracker 405 Blue Mitochondria Dye (Merck #SCT135) using flow cytometry. Per sample 100,000 events were measured using the MACS Quant Analyzer 10 (Miltenyi) and analyzed using the FlowJo_v10.6.1 software. Sexual conversion rates were calculated by dividing the gametocytemia on day 5 by the total parasitemia on day 1 (Extended Data Fig. 7a). To assess gametocyte development and morphology, parasites were induced for sexual commitment as described above and the mFA medium replaced with human serum-containing medium supplemented with 50 mM N-acetylglucosamine (GlcNAc) for six consecutive days with daily medium changes to eliminate asexual parasites^69^. Thereafter, gametocytes were cultured in the absence of GlcNAc. Hemacolor-stained thin blood smears have been prepared every second day to assess gametocyte morphology throughout gametocyte development. On day 9 of gametocyte development, methanol/acetone-fixed thin blood smears were prepared and IFAs performed using rabbit Ab α-Pfg377^70^ and DAPI to determine the gametocyte sex ratios (Extended Data Fig. 7b).

### Immunofluorescence assays

The blood smears from pelleted parasite cultures were prepared and fixed with ice-cold methanol or methanol/acetone (60:40) for 2 min, air dried at RT and stored at -80 °C until further use. Fixed cells were incubated for 1 h in blocking solution (3% BSA in PBS), followed by incubating with primary antibodies for 1 h diluted in blocking buffer (rabbit α-HP1^7^, 1:1,000; mouse IgG1 mAb α-GFP, 1:500, Roche Diagnostics #11814460001; mouse mAb α-Pfs16^68^, 1:500; rabbit α-Pfg377^70^, 1:1,000). Cells were washed three times with PBS and secondary antibody in blocking solution was incubated for 45 min in the dark (for α-GFP and α-Pfs16: Alexa Fluor 488-conjugated α-mouse IgG (Molecular Probes #A11001), 1:500; for α-HP1 and α-Pfs377: Alexa Fluor 568-conjugated α-rabbit IgG (Molecular Probes #A11011), 1:500). Cells were mounted using Vectashield antifade containing DAPI (Vector Laboratories #H-1200).

Microscopy was performed with the Leica Thunder 3D assay fluorescence microscope (for α-Pfg377: 40x objective; for all remaining experiments: 63× objective) equipped with a Leica K5 cMOS camera. The Leica application suite X software (LAS X version 3.7.5.24914) was used by the microscope. Images were processed in Fiji (ImageJ, version 1.53t) using identical settings within each experiment.

### Nuclei isolation for CUT&Tag and nuclear lysates

For CUT&Tag samples, parasite cultures were lightly crosslinked with 0.1% formaldehyde (Sigma, F8775) and incubated for 2 min at 37 °C shaking. Addition of glycine to 0.125 M final concentration was used to stop crosslinking and samples were handled on ice from here onwards. This step was omitted for nuclear lysate preparations for Immunoprecipitation.

Parasites were harvested by 440 g centrifugation for 8 min at 4 °C and washed with ice-cold PBS. Pellet was washed another time with PBS supplemented with 1x EDTA-free Protease Inhibitor (Roche, 04693132001). Parasites were extracted by the addition of saponin (0.05% total concentration) and samples were incubated at room temperature (RT) for 10 min. Nuclei were isolated by carefully transferring the extracted parasite mixture on top of a 0.25 M to 0.1 M sucrose gradient in cell lysis buffer (10 mM Tris pH 8, 3 mM MgCl2, 0.2% NP-40, 1x EDTA free Protease inhibitor (Roche, 04693132001); 15 mL 0.25 M Sucrose and 17.5 mL 0.1M Sucrose for 50 mL tubes, 4 mL 0.25 M Sucrose and 6 mL 0.1 M Sucrose for 15 mL tubes) and centrifuging for 12 min, 3100 g, 4 °C with acceleration and deceleration set to 1 (Eppendorf 5910 Ri, Rotor S-4×400).

Supernatant was carefully removed and nuclei washed in cell lysis buffer (10 min, 3500 g, 4 °C; Heraeus Fresco 21). Afterwards, nuclei were washed with cell lysis buffer containing 20% Glycerol and nuclei pellet was snap frozen in liquid nitrogen prior to storage at -80°C. For CUT&Tag, nuclei counts were obtained using an automatic hemocytometer (BioRad, TC10 Automated Cell Counter) prior to snap freezing.

### GFP pull-down

GFP pulldowns were performed as described previously^71^. Specifically, nuclear pellet was washed with native ChIP digestion buffer (50mM Tris pH7.4, 4mM MgCl_­_, 1mM CaCl_­_, 1x EDTA free Protease inhibitor (Roche, 04693132001)) and resuspended in 4x volume of nuclear extraction high salt buffer (50 mM HEPES-NaOH (pH7.5), 20% glycerol, 420 mM NaCl, 1.5 mM MgCl_­_, 1mM DTT, 2x EDTA free Protease inhibitor (Roche, 04693132001), 0.4% NP40, +2µl TURBO DNase (Invitrogen Cat. #AM2238)). The pellet was dounced (25x; Fisherbrand douncer pellet pestles, Cat. #16339635) and incubated on a roller for 2 h at 4 °C. The lysate was spun down at 17000 g for 10 min and the supernatant was collected. The process was repeat 2 times with 3x pellet volume of nuclear extraction high salt buffer (+TURBO Dnase) and the final lysate volumes (∼10x pellet volume) were pooled. Protein quantification was done with BCA Assay (∼3.2 mg total / 0.8 mg per reaction downstream).

All following steps were carried out on ice or 4 °C. The nuclear extract was diluted to 1.273x volume by adding 0.273x volume of Buffer P* (50 mM HEPES-NaOH (pH 7.5), 18.2% v/v glycerol, 1.5 mM MgCl_­_, 0.93 mM EDTA, 0.4 % NP40, 2x EDTA free Protease inhibitor (Roche, 04693132001), 1 mM DTT). Ethidium Bromide (Sigma E1510) was added to a final conc. of 50 µg/and extract was spun at 17000 g for 25 min to remove freeze-thaw precipitates and supernatant was transferred into a clean tube. lysate pools were split into 4 reactions (2 for GFP-IP and 2 for control-IP). 15 µl of each GFP-Trap® (Chromotek-gta-20) and Blocked Agarose-Beads (BAB/Chromotek-bab-20) were washed 3 times (2000 g, 2 min, 4 °C) with 1 ml Buffer P. The nuclear lysate was then split across two GFP-Trap and two BAB samples, (0.8 mg of extract per IP condition). The reactions were incubated on a rotating wheel for 90 min at 4 °C. Post incubation, the supernatant was collected by centrifugation (2000 g , 2 min). Beads were washed (twice 1 ml Wash Buffer (300 mM NaCl, 50 mM HEPES-NaOH (pH 7.5), 18.2 % v/v glycerol, 1.5 mM MgCl_­_, 0.2 mM EDTA, 0.5 % NP40, 2x EDTA free Protease inhibitor (Roche, 04693132001), 1 mM DTT), twice 1ml PBS/0.5% NP-40, and twice with 1ml PBS. A final wash was given with 100 mM TEAB. Residual buffer was removed with a low bore insulin syringe (BD Micro-fine, Cat. #U-100).Beads were immediately resuspended in 50 µl elution buffer (2M urea, 10 mM DTT, in 100 mM Tris, pH 7.5) and incubated on a shaker for 20 min at 1400 rpm. Iodoacetamide (IAA) (Sigma-Aldrich Cat. #I1149) was added to a final concentration of 50 mM and the reaction was incubated on shaking in dark for 10 min. 2,5 µl sequencing grade modified trypsin (Promega Cat. #V5113; 17184 u/mg stock) was added per tube and incubated on shaker for 2 hr. The samples were spun down and supernatant was collected. Another 50 µl of elution buffer was added to the beads and incubated on the shaker for additional 5 min. Post spin, the second eluate was mixed with the original supernatant fraction and 1 µl of trypsin was added to the mix (∼100 µl) and kept on shaker incubator overnight at room temperature.

C18 membranes were setup in stage-tips and washed with 50 µl methanol, 50 µl Buffer B (80% Acetonitrile/0.1% formic acid in H_­_0) and then 2x with 50 µl Buffer A (0.1% formic acid in H_­_0). Samples were immediately loaded on the equilibrated tips and centrifuged for 10 min at 2400 rpm. Tips were then washed with 50µl Buffer A (600 g for 2 min). A light labelling (3mL dimethyl labeling buffer (10 mM NaH_­_PO_­_, 35 mM Na_­_HPO_­_), 16,2 µl of 37% formaldehyde (CH2O, light, Sigma Cat. #252549) and 6 mg sodium cyanoborohydride (NaBH_­_CN, light, Sigma Cat. #156159) and heavy labelling buffer (3mL dimethyl labeling buffer, 30 µl 20% formaldehyde-d2 (CD2O, heavy deuterium isotope, Sigma Cat. #492620) and 6 mg sodium cyanoborodeuteride (NaBD_­_CN, heavy, Sigma Cat. #205591) were prepared. 300 µl of appropriate labelling buffer was passed through each stage-tip by centrifugation at 2200 g for 5 min. For a single biological replicate set (4 reactions), both GFP and control BAB were labeled with heavy and light label each. A final wash was given with 100 µl Buffer A, spinning at 1500 g for 5 min. The samples were then eluted from the stage tip and loaded onto the mass spectrometer (1hr run on Exploris).

### Mass-spec data analysis in MaxQuant

Data analysis of raw mass spectrometry files was done on the MaxQuant software (v2.1.4). The forward and reverse experiments were combined from the two biological replicates. The software was configured as per the following parameters: Variable modifications (deamidation (NQ), oxidation (M), Acetyl (protein N-term)), fixed modification (carbamidomethyl (C)) with max 5 modifications per peptide. Digestion was specified for Trypsin/P. The run was configured for re-quantification. The ratios of H/L normalized forward and reverse experiments were plotted using a custom R script for identifying significantly enriched proteins in pull-down.

### CUT&Tag reaction & library preparation

We have adapted the protocol of based on Kaya-Okur et. al ^72^ to malaria parasites with changes regarding permeabilization; DNA extraction; Library preparation protocol as already described in our previous work^45^. Specifically, 1 million isolated nuclei were used as input material for all reactions. For primary antibody incubation, 0.25 µL polyclonal rabbit αHP1 ^7^in 50 µL antibody buffer (20 mM HEPES pH 7.5, 150 mM NaCl, 0,5 mM Spermidine, 2 mM EDTA, 0.1% BSA, 1x EDTA-free protease inhibitor) was added and incubated nutating over night at 4 °C. Secondary antibody incubation (1.2 µg guinea pig anti-rabbit antibody, Antibodies-Online ABIN101961) in 100 µL CUT&Tag wash buffer (20 mM HEPES pH 7.5, 150 mM NaCl, 0,5 mM Spermidine, 1x EDTA-free protease inhibitor), was performed at a 1:100 dilution for 1 h on RT, nutating. 2.5 µL commercial proteinA/G-Tn5 fusion protein (CUTANA™ pAG-Tn5 for CUT&Tag, Epicypher, 15-1017) was added in 50 µL CUT&Tag 300 wash buffer (20 mM HEPES pH 7.5, 300 mM NaCl, 0,5 mM Spermidine, 1x EDTA-free protease inhibitor) and incubated nutating for 1 h. The supernatant was removed and the nuclei were resuspended in 200 µL freshly prepared tagmentation buffer (CUT&Tag 300 wash buffer, 10 mM MgCl_­_). To perform tagmentation, samples were incubated in a PCR thermocycler (BioRad, T100) at 37 °C for 1 h. Tagmentation was stopped and nuclei or parasite lysis was facilitated by addition of 10 µL of 0.5M EDTA pH8, 3 µL of 10% SDS and 1 µL of 50 mg/mL proteinase K. Samples were briefly vortexed and then incubated at 55 °C for 1 h. DNA fragments were extracted utilizing the DNA Clean & Concentrator - 5 kit (Zymogen, D4014) following manufactures instructions. DNA was eluted from the column with 26 µL of prewarmed elution buffer and DNA concentrations were assessed with Qubit dsDNA High Sensitivity Assay kit (Invitrogen, Q33231). Maximal 50 ng of extracted DNA from CUT&Tag experiments were amplified with unique combinations of i5 and i7 barcoded primers ^73^. PCRs were performed in a total reaction volume of 50 µL using Kapa HiFi polymerase (use non-hotstart version for gap filling; Roche, Cat# 07958838001) with the following program: 58 °C for 5 min, 62 °C for 5 min (gap filling), 98 °C for 2 min, 12-16 cycles of 98 °C for 20 second and 62 °C for 10 seconds, 62 °C for 1 min and hold at 4 °C. Post-PCR DNA cleanup was performed by adding 50 µL (1x volume) of AMPure XP bead slurry (Beckman Coulter, A63882) and incubating for 10 min at RT, washing twice with 80% EtOH on a magnetic rack, and eluting in 16.5 µL of 10 mM Tris-HCl pH 8 for 5 min at RT. DNA concentrations of libraries were assessed with Qubit dsDNA High Sensitivity Assay kit (Invitrogen, Q33231) and library fragment size distribution was accessed by microfluidic gel electrophoresis (Agilent 2100 Bioanalyser) with the corresponding High Sensitivity DNA Kit (Agilent, 5067-4626). Sequencing was performed using an Illumina NextSeq 2000 instrument for 10 M reads per CUT&Tag sample; 59bp paired-end reads were generated.

### CUT&Tag Data analysis

Sequencing reads were mapped against the reference genome PlasmoDB v26 3D7 using bowtie2 (v2.5.2)^74,75^ with paired-end mapping. Duplicate removal was skipped for CUT&Tag datasets as duplicates may result due to the affinity of Tn5 to certain sequences as well as accessibility of certain regions leading to fragments with the same start and end locations^76^.

Reads were filtered by mapping quality >= 30 and mitrochondrial as well as apicoplast reads were removed with samtools (v1.18)^77^. BigWig files normalized to read per million per kilobase (RPMK) were created using deeptools (v3.5.4)^78^. BedGraph files were generated for visualisation purposes on the UCSC genome browser^79^ with bedtools (v2.31.0)^80^ normalized to library size. A detailed & customisable script can be found on our github page https://github.com/bartfai-lab/DiBio-CUTnTag-Analysis or 10.5281/zenodo.15638929.

Chromosomal genome-wide overview (Extended Data Fig 4) was generated by exporting visualized tracks from the UCSC genome browser, compiling and scaling them to each other in Adobe Illustrator.

Bedgraph files of replicates were combined into one file with bedtools unionbedg (v2.31.0)^80^, averages per position calculated and a new bedgraph file generated.

### RNA harvest and isolation

3D7 wild type and 3D7/JmjC1/2-dKO parasites were tightly synchronized to a ∼4 h window using repeated sorbitol treatments and total RNA samples were harvested at 10-14, 20-24, 30-34 and 40-44 hpi in biological triplicates. Giemsa-stained thin blood-smears prepared for each sample taken to assess stage progression and synchronicity between the cultures and replicates. 10 mL culture aliquots (5% hematocrit, >5% parasitemia) were harvested for the 10-14 and 20-24 hpi samples, 5 mL for 30-34 and 40-44 hpi samples. For harvesting, cultures were spun down (440 g, 5 min) and the supernatants removed. The RBC pellets were resuspended in 0.5x of aliquot volume of Trizol, pellets were resuspended and samples were stored at -80 °C.

For RNA isolation, total RNA was purified using the Direct-zol RNA Miniprep Kit (ZYMO #R2050). Samples were thawed and transferred into a fresh tube to limit carry-over of debris. RNA isolation with on-column DNA digestion was performed according to manufacturer’s instructions. RNA was eluted twice with 50 µL nuclease-free water. To perform a second round of DNaseI treatment to minimize DNA contamination, 350 µL RLT buffer (QIAGEN, Cat# 79216) was added to the elutions and 250 µL freshly prepared 70% EtOH was added and carefully mixed. The mixture was applied to a new Direct-zol column and the DNaseI incubation was repeated, followed by washes with 400 µL RNA prewash buffer and 700 µL RNA wash buffer. RNA was eluted twice in separate fractions E1 and E2 with 30 µL nuclease-free water each. 5-10 µL aliquots were prepared and frozen at -80 °C. RNA concentration was measured using Qubit RNA High Sensitivity Kit (Thermo Fisher Scientific, #Q32855) and integrity of RNA was assessed by microfluidic gel electrophoresis (Agilent 2100 Bioanalyser) with the RNA 6000 Nano kit (Agilent, 5067-1511).

### RNA sequencing library preparation

250 ng of RNA were used to generate libraries with the KAPA HyperPrep Kit according to manufactures instructions with the following adjustments: Step 4 (A-tailing) was performed at 55 °C for 20 min. Library amplification PCR protocol was adapted to 98 °C for 2 min, 12 cycles of 98 °C for 20 s and 62 °C for 2 min, final extension for 2 min at 62 °C and hold at 4 °C. Sequencing was performed using an Illumina NextSeq 2000 instrument for 20 M reads per RNA-seq sample; 59bp paired-end reads were generated.

### RNA-seq data analysis

A detailed and customisable script for the RNAseq mapping can be found at our github page: https://github.com/bartfai-lab/JmjCs_heterochromatin-calling_RNAseq_P.falciparum

Quality of demultiplexed sequencing files was checked by fastqc (v0.12.1) and adapters trimmed using trimgalore (v0.6.10) with the parameters -q 30, -j 8 with the paired flag. Indexes were generated using STAR (v2.7.11b), and trimmed reads were mapped to the PlasmoDB v68 3D7 reference genome using the STAR aligner with --quantMode GeneCounts. Unique alignments were filtered with samtools (v1.21) view -q 255. Duplicate alignments were removed by samtools markdup and reads counted using featureCounts from the subread package (v1.5.3), generating a count matrix.

Count matrix was further analysed in R using the DESeq2 package (1.42.1) to obtain differential expressed genes. Gene filters were set to retain only genes which had at least 10 reads in either WT or KO condition and DESeq2 analysis was done separately for each time point. Significant differential expression was defined as log2 expression fold change of > 2 (upregulated or < -2 (downregulated) and Benjamini-Hochberg adjusted *P*-value < 0.01. PCA was performed to check for potential batch effects of replicates and WT / KO growth delays (Extended Data Fig. 4a).

Clustering was performed on k-means = 7 using morpheus^81^, and heatmaps were plotted in R using the pheatmap package (v 1.0.13).

Additionally to DESeq2 analysis, TPM expression values for each replicate and timepoint (mean and standard deviation) were calculated in R.

### Micro-C

The Micro-C protocol was performed as previously described^50^ (Singh et al., 2025). 3D7 wild type and 3D7/JmjC1/2-dKO parasites were synchronized, and eight replicates (>5% parasitemia, 1 mL blood at 5% hematocrit) were collected at 40-44 hpi.

Prior to harvest, leucocytes were removed from cultures by filtration using a Plasmodipur filters (Europroxima 8011) according to manufactures instructions. A single filter was used per 5 mL of packed iRBC diluted in ∼40 mL media. After filtration, parasites were centrifuged and lysed with saponin (0.075% in DPBS) and washed with DPBS at 37 °C. Parasites were resuspended in DPBS at 25 °C and cross-linked for 10 min by adding methanol-free formaldehyde (ThermoFisher 28908) to 1% final concentration with gentle agitation. The reaction was quenched by adding 1 M Tris-HCl pH 7.5 to a final concentration of 0.75 M and incubating at 25 °C for 5 min with gentle agitation. Parasites were centrifuged for 5 min at 3250 g, washed with DPBS at 25 °C, and crosslinked a second time with 3 mM DSG (ThermoFisher 20593) in DPBS for 45 min at 25 °C with agitation. The reaction was quenched with 1 M Tris-HCl pH 7.5 to a final concentration of 0.75 M and incubated at 25 °C for 5 min with agitation. The double cross-linked parasites were washed with DPBS at 25 °C, and the pellets were snap-frozen and stored at -80 °C until further use.

One replicate was split into four tubes and used for MNase titration. Each pellet was resuspended in 1 mL of MB#1 [50 mM NaCl, 10 mM Tris-HCl pH 7.5, 5 mM MgCl_­_, 1 mM CaCl_­_, 0.2% NP-40, 1x protease inhibitor (PI, Roche 4693159001)] and incubated on ice for 20 min with regular flicking. The cells were centrifuged (10000 g, 5 min, 4 °C), washed with MB#1, resuspended in 760 μL of MB#1, and aliquoted in 4 tubes of 190 μL. 10 μL of MNase (Worthington Biochem LS004798) at different concentrations (for titration) was added, and the mixture was incubated for exactly 10 min at 37 °C with continuous shaking at 850 rpm. To stop the reaction, 500 mM EGTA (ThermoFisher J60767) was added to a final concentration of 4 mM and was followed by a 10 min incubation at 65 °C. The digested nuclei were then washed twice (5 min centrifugation at 16000 g, 4 °C) with MB#2 (50 mM NaCl, 10 mM Tris-HCl pH 7.5, 10 mM MgCl_­_). The nuclei were resuspended in a digestion mix (10 mM Tris-HCl pH 7.5, 1 mM EDTA, 1% SDS, 1 mg/mL Proteinase K (ThermoFisher EO0491), 0.125 mg/mL RNase A (ThermoFisher EO0491)) and de-crosslinked for 10 h at 65 °C. The samples were centrifuged (16000 g, 10 min, 4 °C) and DNA contained in the supernatant was extracted using the DNA Clean & Concentrator kit (Zymo D4013). The fragment sizes were assessed by microfluidic gel electrophoresis (Agilent 2100 Bioanalyser) with the corresponding High Sensitivity DNA Kit (Agilent, 5067-4626) to choose the appropriate MNase concentration yielding a 90/10 monomer/dimer ratio (250U for 3D7 WT and 25U for JmjC1/2 KO nuclei pellets).

The following steps were performed with four replicates for late-stage parasites, respectively. The nuclei pellet was lysed and digested with the chosen concentration of MNase, as described above. After the MB#2 washes, each pellet was centrifuged (16000 g, 5 min, 4 °C), resuspended in 45 μL of end-chewing mix (1X NEB Buffer 2.1, 2 mM ATP (ThermoFisher R1441), 5 mM DTT (ThermoFisher 707265ML), 0.5 U/μL T4 PNK (NEB M0201S)) and incubated at 37 °C for 15 min on an agitator with interval mixing (800 rpm 10 s, resting 30 s). 5 μL of 5 U/μL Klenow fragment (NEB M0210L) was added and incubated at 37 °C for 15 min with the same agitation program as the previous step. 25 μL of end-labeling mix (0.2 mM Biotin-dATP (Jena Bioscience NU-835-BIO14-S), 0.2 mM Biotin-dCTP (Jena Bioscience NU-809-BIOX-S), 0.2 mM dTTP (ThermoFisher 18255018), 0.2 mM dGTP (ThermoFisher 18254011), 0.1 mg/mL BSA (Invitrogen AM2616)) was added to the mixture and incubated at 25 °C for 45 min with agitation (800 rpm 10 s, resting 30 s). 500 mM EDTA (Invitrogen AM9260G) was added to a final concentration of 30 mM and incubated for 30 min at 65 °C. The pellet was centrifuged (10000 g, 5 min, 4 °C) and washed with cold MB#3 (10 mM Tris-HCl pH 7.5, 10 mM MgCl_­_). The biotinylated

DNA fragments were then covalently linked by proximity ligation by resuspending the pellet in 250 μL of ligation mix (1X T4 DNA ligase buffer (NEB B0202S), 0.1 mg/mL BSA (Invitrogen AM2616), 33 U/μL T4 DNA ligase (NEB M0202L)) and incubating at 25 °C for 2.5 h with slow rotation. The ligated DNA fragments were then centrifuged (16000 g, 5 min, 4 °C), and the biotin-dNTPs were removed from non-ligated ends by resuspending the pellet in 100 μL of 1x NEB buffer #1 (NEB B7001S) and 5 U/μL exonuclease III (NEB M0206L) and incubating for 15 min at 37 °C with agitation (800 rpm 10 s, resting 30 s).

After addition of 12.5 μL of 10% SDS (ThermoFisher 15553027) DNA was de-crosslinked by incubating for 10 h at 65 °C. RNase A (ThermoFisher EN0531) was added to a final concentration of 0.2 mg/mL and incubated for 2 h at 37 °C. Proteinase K (ThermoFisher EO0491) was added to a final concentration of 0.2 mg/mL and incubated for 2 h at 55 °C. After centrifugation (16000 g, 10 min, 4 °C), the DNA contained in the supernatant was recovered using the DNA Clean & Concentrator kit (Zymo D4013) using a 5:1 ratio of DNA Binding Buffer to sample. The ligated DNA was mixed with loading dye (ThermoFisher R1161) and loaded onto a 2% TAE agarose gel to separate dimers from monomers. The gel was run at 100 V until a satisfying separation was obtained. A band from 250 - 400bp was cut and DNA was extracted in 150 μL of ddH2O using the Gel DNA Recovery Kit (Zymo D4007).

The biotinylated dimers were isolated with Dynabeads MyOne Streptavidin C1 (ThermoFisher 65001). The beads were washed with 500 μL TBW (1 M NaCl, 5 mM Tris-HCl pH 7.5, 0.5 mM EDTA, 0.10% Tween20) on a magnet and resuspended in 150 μL 2X BW (2 M NaCl, 10 mM Tris-HCl pH 7.5, 1 mM EDTA). The ligated fragments were added and rotated for 20 min at 25 °C. The beads were then washed twice with 500 μL TBW at 55 °C on an agitator (2 min, 800 rpm for 10 s, resting 30 s) and once with EB (10 mM Tris-HCl pH 7.5) at 25 °C. The library prep was performed with the samples on beads using the NEBNext® Ultra II Directional DNA Library Prep Kit (NEB E7645), with several modifications. First, the end repair and adaptor ligations steps were performed with interval mixing (800 rpm for 10 s, resting 30 s). After the adaptor ligation, the beads were washed twice with TWB at 55 °C and once with EB (as described above) and resuspended in 20 μL EB for the PCR reaction. No size-selection step was performed. Finally, after the PCR, the beads were placed again on a magnet, and the supernatant was purified with AMPure XP beads as recommended.

### Micro-C computational processing and analysis

Micro-C paired-end reads (150 bp paired end) were mapped to the *P. falciparum* genome (plasmoDB.org, version 3, release 56) and processed into pairs and multi-resolution normalized contact matrix.mcool files using hicstuff (v3.2.2) (https://github.com/koszullab/hicstuff, ^82^) with the following non-default options: ‘--enzyme mnase --mapping iterative --duplicates --binning 100’. Individual replicates pairs files were subsequently merged using pairtools^83^ and multi-resolution binned contact matrix files (.mcool files) of merged replicates were regenerated using cooler (v0.9.3)^84^. When comparing control to 3D7/JmjC1/2-dKO Micro-C results, 3D7/JmjC1/2-dKO contacts were subsampled with pairtools (v1.0.3) to match 3D7 wild type Micro-C data.

Observed/expected contact frequency was computed using the ‘detrend()’ function from HiContacts v 1.8.0) ^85,86^. Aggregated contact maps were generated using the ‘aggregate()’ function from HiContacts. All contact maps, including aggregate plots, were generated in R using HiContacts. Cut & Tag generated HP1 regions and resulting heterochromatin gain, and loss regions were used for identifying heterochromatin interactions and construct aggregate contact maps.

Long-range interactions were annotated in Micro-C data using the automated structural feature caller chromosight (v1.6.3), using 2 kb resolution for long-range interactions and filtering for q-values ≤ 10-3. Visualization of cis interactions were done using IGV (v2.18.2). The 1D aggregate HP1 tracks were plotted using tidyCoverage in R^87^.

### Protein sequence analysis

PfJmjC1 domains were annotated using InterPro on the PlasmoDB database^44^. Structural similarity of PfJmjC1 to human KDM5A was identified by similarities to protein data bank sequences on PlasmoDB. Different domain structures of Jumonji-C domain containing protein families as described by Klose et al^32^ were compared to PfJmjC1 and selected proteins from different species aligned using Clustal Omega MSA^88^. Multisequence alignment was visualized using Jalview (v2.11.4.1). Binding residues for Fe (II) Ions and αKG were marked according to Klose et al^32^.

### Identification of heterochromatic regions

Heterochromatin domains were identified utilizing a custom script, which can be found on our github page: https://github.com/bartfai-lab/JmjCs_heterochromatin-calling_RNAseq_P.falciparum

In short, average enrichment scores of previously generated bigwig files from CUT&Tag data were generated using multiBigwigSummary from the deep tools package. Bins were filtered on enrichment values (filter = 100 tags in a 500bp bin, chosen based on distribution of total avg enrichment values). Neighboring bins with enrichment were then combined using bedtools merge, allowing for skipping of filtered bins depending on a predetermined binskip value (x*binsize, x=20 used for analysis). Lastly, only bins of a minimal size were kept analysis (minimal bins (minBin), x*binsize , x=10 used for analysis). Final heterochromatin calls were saved as a bed file and metadata such as filter and annotated bp were saved in a txt file.

### Identification of *de novo* heterochromatinised regions

Replicate average bedgraphs were sorted and transformed into bigwig files using bedGraphToBigWig. Log2 ratios between 3D7 WT and 3D7/JmjC1/2-dKO mutant were calculated using a custom script, which can be found on our github page (https://github.com/bartfai-lab/JmjCs_heterochromatin-calling_RNAseq_P.falciparum). In essence, average enrichment over 100 bp bins was calculated utilising multiBigWigsummary. Apicoplast and mitochondrial regions were removed and regions were filtered for a minimum enrichment score of 3 in either bigwig files. This step reduces noise due to sparce sequencing data and high foldchanges between regions at areas of low coverage. A count of 1 is added to all remaining regions to avoid dividing by 0, and log2 ratios are calculated. Bedgraphs are generated and further transformed into BigWig files with bedGraphToBigWig.

For *de novo* peak analysis, log2 ratio BigWig files are subjected to our heterochromatin calling script to identify greater than 2 fold increases and decreases in comparative heterochromatin occupancy using a filter of 1 or -1 respectively with a binskip value of 20 and minBin value of 10. This resulted in the identification of 63 *de novo* heterochromatin regions in the 3D7/JmjC1/2-dKO (Extended Data Table 1). 3D7/JmjC1/2-dKO -specific peaks were identified by calculating 3D7/JmjC1-KO / 3D7 wild type enrichment values in these 63 regions using multiBigWigsummary. Regions were classified as 3D7/JmjC1/2-dKO specific *de novo* regions if values were lower or equal than 0.7 (log2), corresponding a ∼1.6 fold change towards wildtype. This cutoff was chosen to accommodate smaller increases in heterochromatin in the 3D7/JmjC1-KO level.

Focusing on regions of heterochromatin loss, we discovered that the Chromosome 14 distal end has most likely been lost in a sub population of our 3D7/JmjC1/2-dKO parasites (Extended Data Fig. 5e). This is further indicated by an apparent lack of reads mapping to this region in our RNAseq data, and we therefore excluded this chromosome end from our analysis.

For enrichment analysis over gene bodies or regulatory regions, multiBigWigsummary was used to calculate average enrichment of log2 ratios of 3D7/JmjC1/2-dKO and 3D7 wild type in these areas utilising bed files for either gene body or proposed regulatory region (-1000bp ATG +500bp) locations. Results were filtered for areas with enrichment values greater than 1 (log2), corresponding to a 2-fold increase of signal in the 3D7/JmjC1/2-dKO compared to wild type. For maximum expression analysis, gene IDs were identified and corresponding transcriptomics data identified from PlasmoDB utilising the Toenhake et al dataset for asexual expression^51^, the Lasonder et al dataset for female / male gametocyte expression^89^ and the Lopez-Baragan dataset for ookinete expression^90^.

Occupancy profiles were plotted using the computeMatrix function in the deeptools package with the following settings: scale-regions, --binSize 100, --beforeRegionStartLength 5000, --regionBodyLength 5000, --afterRegionStartLength 5000, --skipZeros. Three replicate averaged BigWigs were plotted over locations of genes with increased heterochromatin occupancy using plotProfile and resulting profiles were merged into one panel using Adobe Illustrator.

### GC-content calculation

Random euchromatic and heterochromatic regions as defined by 3D7 wild type HP1 CUT&Tag were selected with the same sizes as 3D7/JmjC1/2-dKO peak regions excluding 3D7 Wild type heterochromatic regions using bedtools shuffle (v2.31.0)^80^ for 10 iterations and resulting bed files merged. GC content was calculated for both generated euchromatic background and 3D7/JmjC1/2-dKO *de novo* heterochromatin locations using bedtools nuc (v2.31.0)^80^. Violin plots were generated using the ggplot2 package (v3.4.4)^91^and Wilcoxon signed-rank test was carried out in R.

### *de novo* motif search

*De novo* motif search was conducted using the HOMER package (v5.1) and JmjC1/2-dKO specific regions as annotated in Extended Data Table 1 running the findMotifsGenome.pl command with -size 200 and 1000, respectively. Top 5 *de novo* results are shown in Extended Data Fig. 5a, though no convincing motif was identified.

## Extended Data

**Extended Data Fig. 1.**
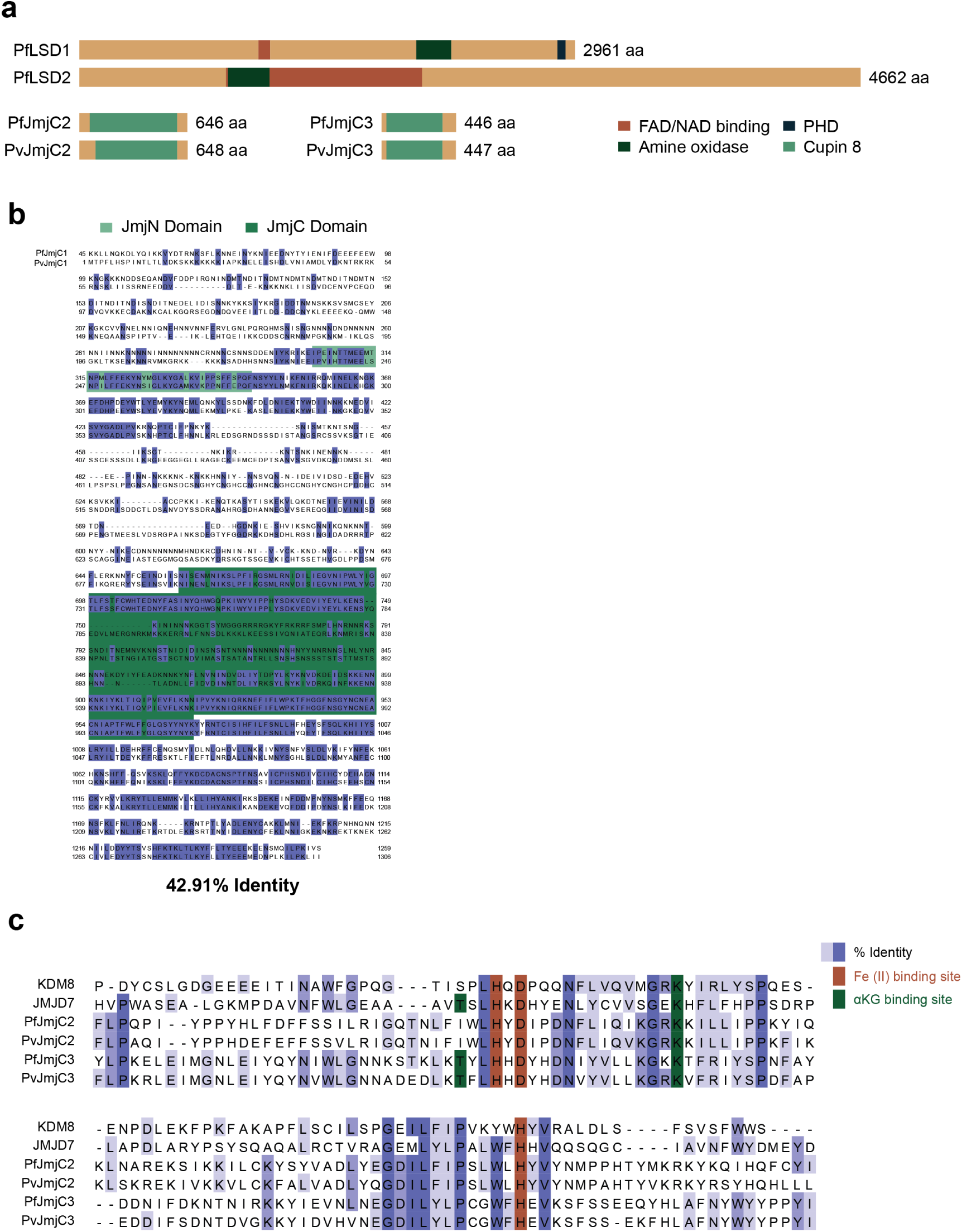
**a)** Domain structure of P. falciparum LSDs as well as P. falciparum and P. vivax JmjC2 and JmjC3. Domains were annotated according to InterProtScan. **b)** Pairwise alignment of PfJmjC1 and PvJmjC1. JmjN (light green) and JmjC (dark green) are marked and show high amino acid conservation. **c)** Multiple sequence alignment of the JmjC domains of proteins related to PfJmjC2 and PfJmJC3, highlighting two separate parts of the JmjC domains with co-factor binding sites. Fe(II) binding sites are marked in red, αKG binding sites in green and level of amino acid (aa) identity with a blue hue.

**Extended Data Fig. 2.**
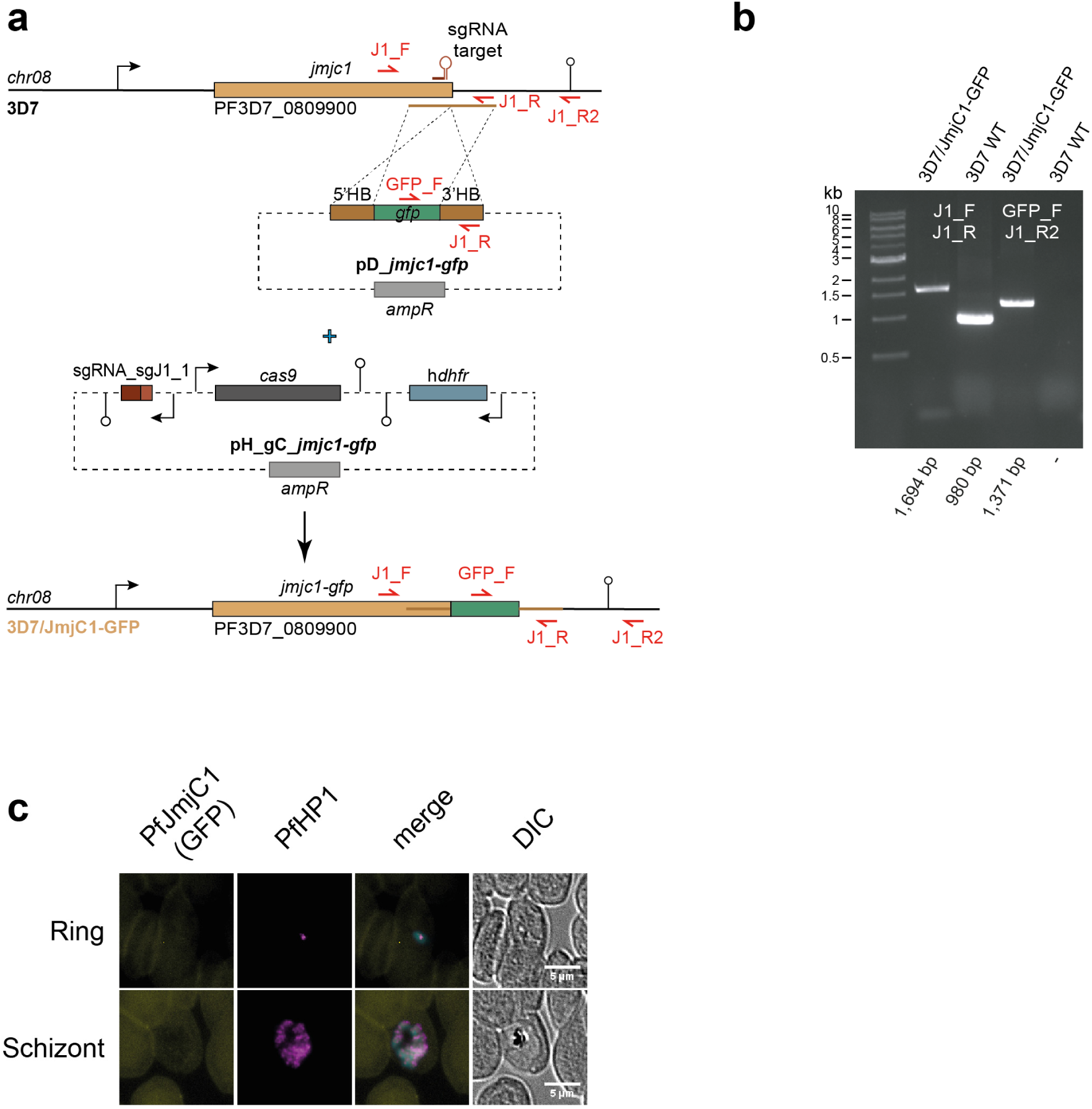
**a)** Schematic illustration of the wild type (top) and modified (bottom) pfjmjc1 locus, along with the pD_jmjc1-gfp donor and pH_gC-jmjc1-gfp CRISPR/Cas9 plasmids used to engineer the 3D7/JmjC1-GFP parasite line. Names and relative positions of primers used for diagnostic PCRs are indicated in red. Note that the three introns of the pfjmjc1 gene are not depicted. HB, homology box. **b)** PCRs performed on genomic DNA extracted from the transgenic 3D7/JmjC1-GFP line and 3D7 wild type (WT) parasites as control. Names of primer pairs used for diagnostic PCRs are indicated in white. Expected DNA fragment sizes are given below the gel images. kb, kilo base. **c)** Representative examples of IFAs of PfJmjC1-GFP and PfHP1 in the 3D7/ PfJmjC1-GFP parasite line at ring and schizont stages. Anti-GFP signals are shown in yellow and anti-PfHP1 in magenta. DAPI was used to stain the parasite genome (blue).

**Extended Data Fig. 3.**
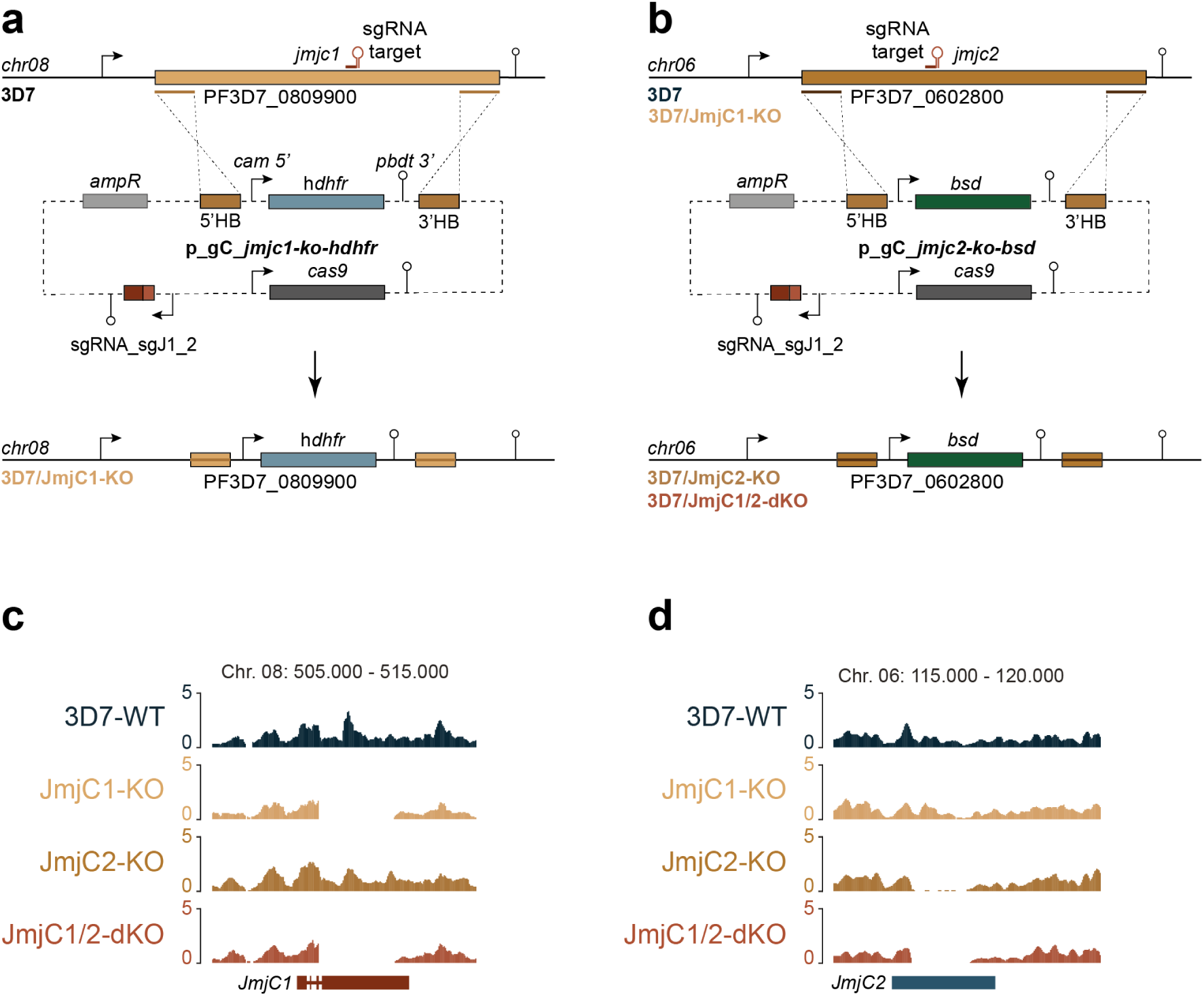
**a)** Schematic illustration of the wild type (top) and modified (bottom) pfjmjc1 locus, along with the p_gC-jmjc1-ko-hdhfr CRISPR/Cas9 all-in-one plasmid used to engineer the 3D7/JmjC1-KO parasite line. Note that the three introns of the pfjmjc1 gene are not depicted. **b)** Schematic illustration of the wild type (top) and modified (bottom) pfjmjc2 locus, along with the p_gC-jmjc2-ko-bsd CRISPR/Cas9 all-in-one plasmid used to engineer the 3D7/JmjC2-KO and 3D7/JmjC1/2-dKO parasite lines. HB, homology box. hdhfr, human dihydrofolate reductase gene; bsd, blasticidin deaminase; cam 5’, P. falciparum calmodulin gene promoter; pbdt 3’, P. berghei dhfr-ts terminator.JmjC1-KO and b) JmjC2-KO. **c,d)** Validation of pfjmjc1 (c) and pfjmjc2 (d) gene deletions in the three KO mutants utilising PfHP1 CUT&Tag occupancy tracks. Deleted regions do not have mapped reads as expected from each mutant.

**Extended Data Fig. 4.**
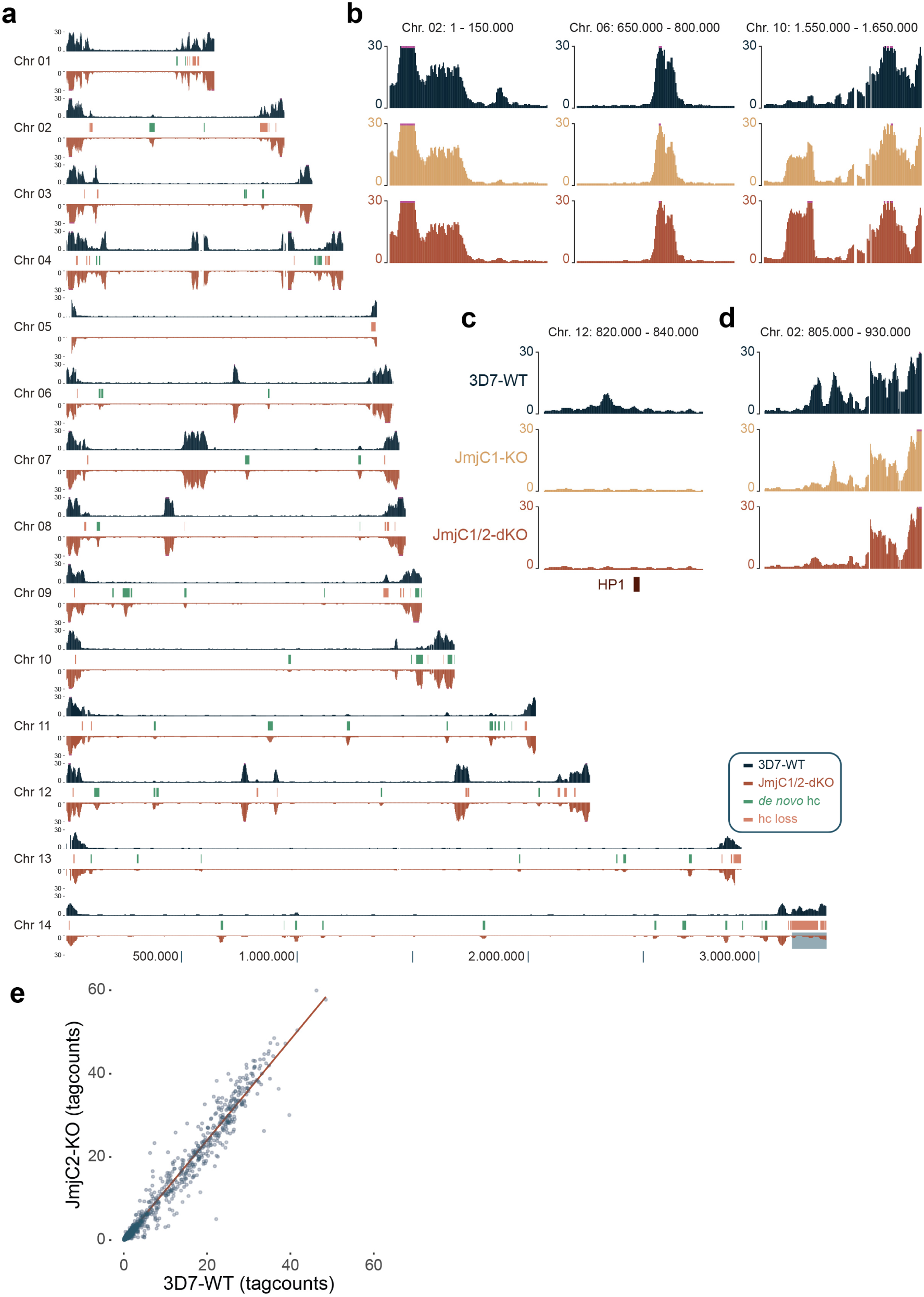
**a)** PfHP1-occupancy tracks for all chromosomes for 3D7 wild type (WT) (dark blue) and 3D7/JmjC1/2-dKO parasites (dark red). Regions identified as de novo heterochromatinised are marked in light green, areas of heterochromatin loss in red. The distal end of chr 14 is lost in the mutant and hence indicated with a blue hue. **b,c,d)** Read occupancy tracks from PfHP1 CUT&Tag experiments, averages of three replicates. (b) Representative example of maintained heterochromatin boundaries in 3D7 wild type (WT), 3D7/JmjC1-KO and 3D7/JmjC1/2-dKO parasites, (c) HP1 locus, (d) representative example of heterochromatin loss at subtelomeric region. **e)** Scatterplot displaying genome-wide correlation of HP1 occupancy (CUT&Tag tagcount) between 3D7 wild type (WT) JmjC2-KO in coding regions.

**Extended Data Fig. 5.**
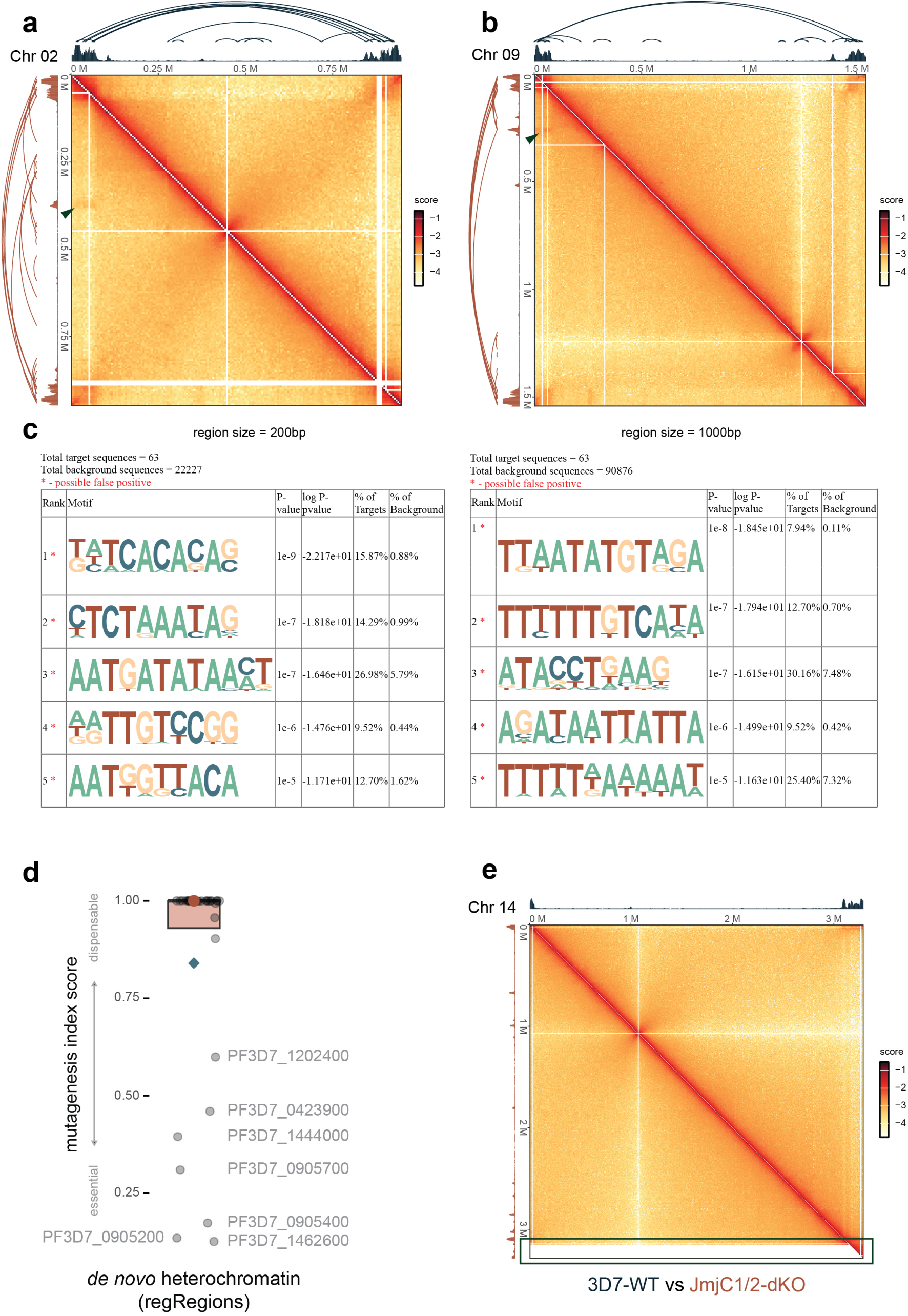
**a,b)** Micro-C contact map of chromosome 2 (a) and chromosome 9 (b) in 3D7 wild type (top right) and PfJmjC1/2-dKO (bottom left) parasites (5kb resolution, normalized interaction frequency) generated from four separate replicates. Color scales are shown at the right. PfHP1 CUT&Tag tracks and identified interactions for 3D7 wild type (dark blue) and 3D7/JmjC1/2-dKO (red) parasites are displayed on the left side and on top, respectively. Strain-specific interactions in the 3D7/JmjC1/2-dKO and 3D7 wild type (WT) parasites are marked with green arrowheads. **c)** Top 5 hits of de novo motif search by the HOMER algorithm, using the sequence of de novo heterochromatin regions in the 3D7/JmjC1/2-dKO parasite strain with and two different region sizes being scanned (200 and 1000bp. All shown motifs are predicted to be false positives and do not show convincing enrichment in de novo regions compared to background. **d)** Boxplot showing mutagenesis index score of genes with de novo heterochromatin at their regulatory regions. Boxplots show quartiles 1 to 3, median as red dot and mean of all data points as blue diamond. Mutagenesis index scores close to 0 mean that the gene is essential, while scores close to 1 mean that the gene is dispensable. Scores taken from Zhang et. al 2018^38^. **e)** Micro-C contact map of chromosome 14 in 3D7 wild type (top right) and PfJmjC1/2 double KO (bottom left) parasites (5kb resolution, normalized interaction frequency) generated from four separate replicates. Color scales are shown at the right. PfHP1 CUT&Tag tracks for 3D7 wild type (dark blue) and 3D7/JmjC1/2-dKO (red) are displayed on the left side and on top, respectively. Potential chromosome end loss in the 3D7/JmjC1/2-dKO is indicated by the lack of interactions and is marked. with a green box.

**Extended Data Fig. 6.**
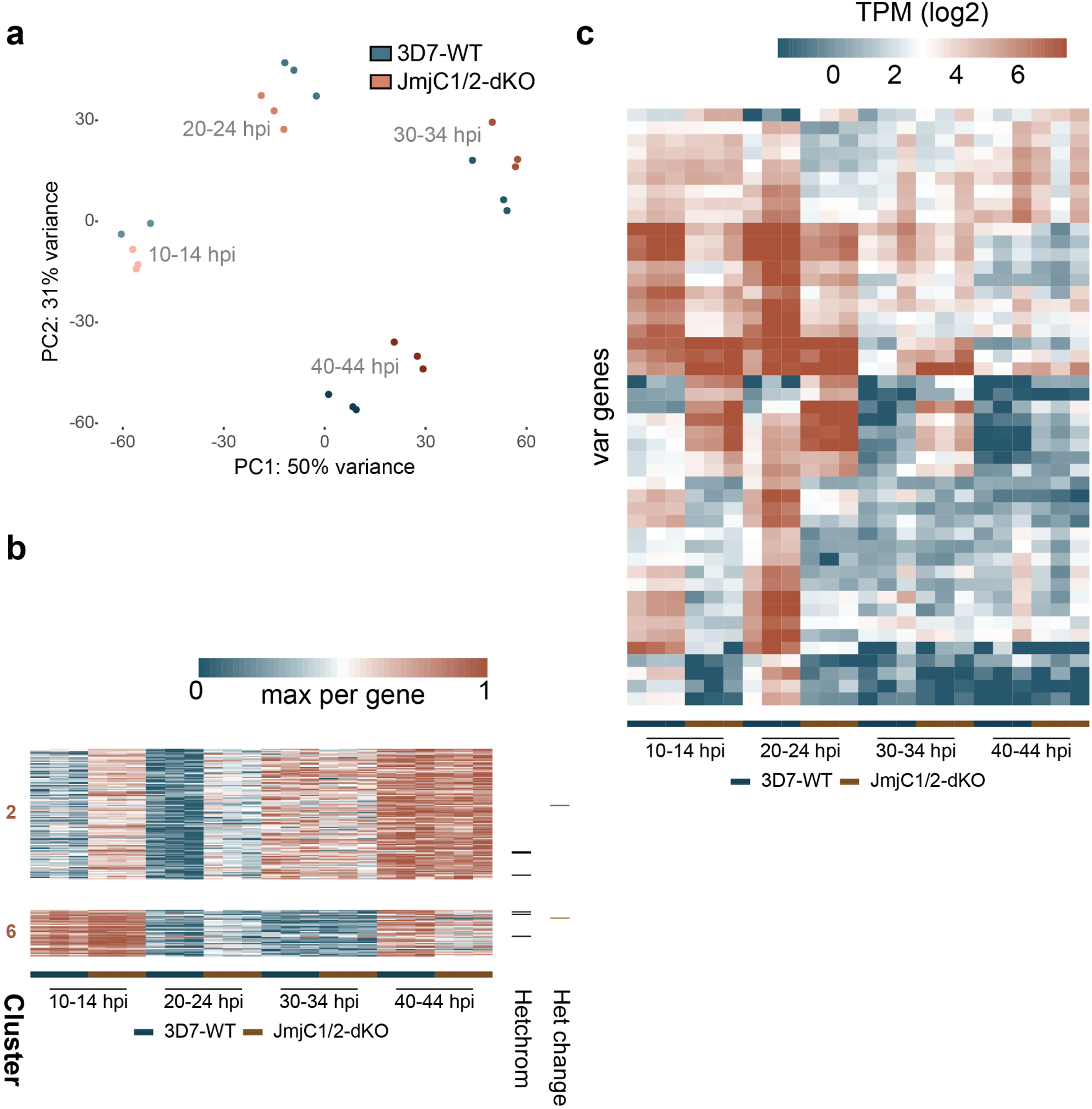
**a)** PCA of vsd transformed values generated by DESeq2 from RNA-seq data off all timepoints of 3D7 wild type (WT) (blue) and PfJmjC1/2-dKO parasites (red). The clear clustering of replicates as well as mutant and WT samples indicate consistency and the lack of substantial developmental delay in mutant parasites. **b)** Heatmap of RNA-seq data of genes differentially expressed between 3D7 wild type (WT) and 3D7/JmjC1/2-dKO parasites. Gene expression values represent three replicate experiments per condition / timepoint and have been scaled to their maximum value per gene. Clustering was performed with k-means and heterochromatin state in wild type parasites is indicated by a black bar (Fraschka et. al, 2018)^8^. Changes in heterochromatin occupancy comparing 3D7 wild type (WT) to 3D7/JmjC1/2-dKO parasites are shown as green (heterochromatin-depleted) or brown (heterochromatin-gaining) bars. Only cluster 2 and 6 are depicted, other clusters are displayed on Figure 4a. **c)** Heatmap of RNA-seq data in log2(TPM) from all differentially expressed var genes.

**Extended Data Fig 7:**
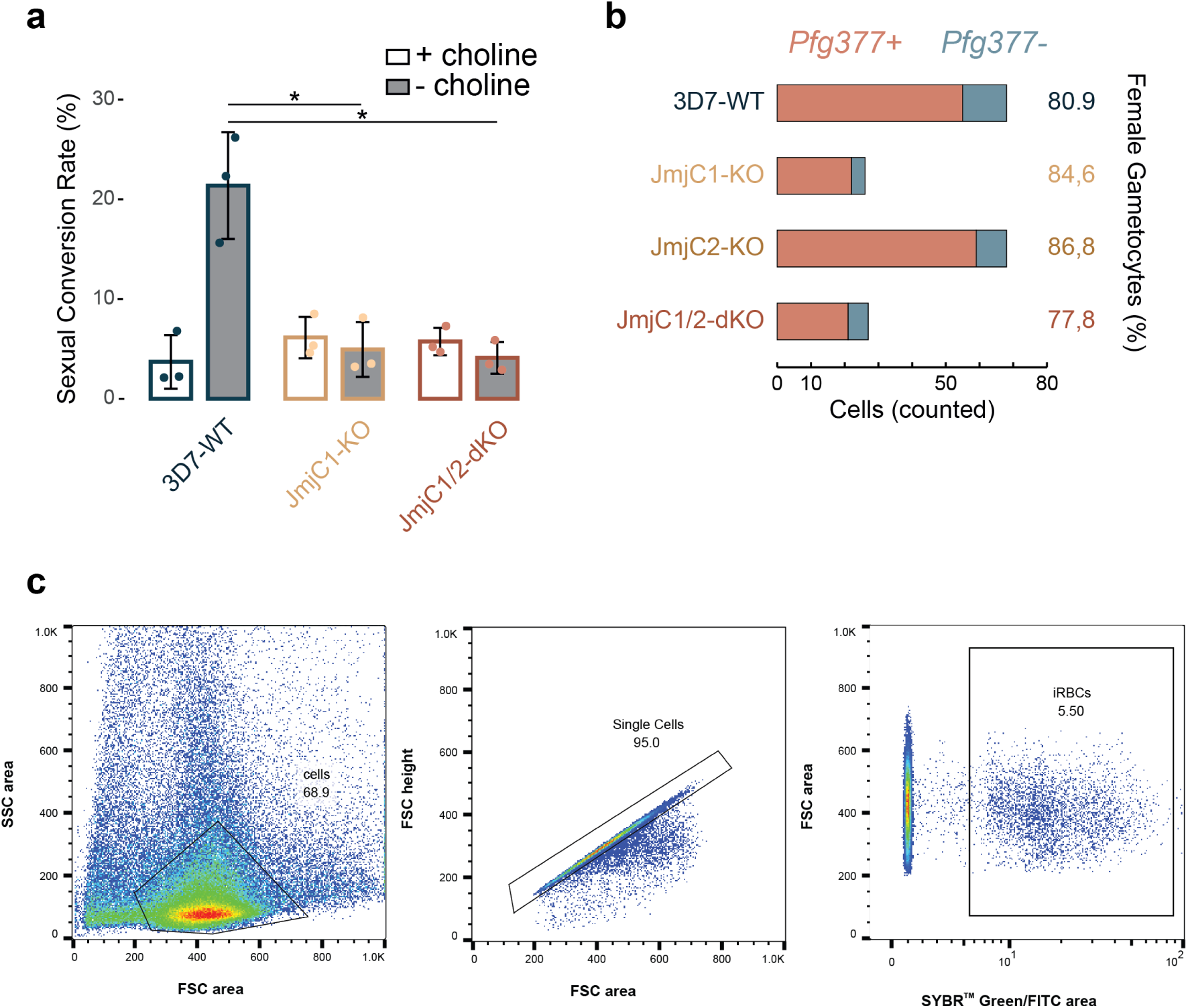
Gating strategy of flow cytometry data obtained from parasite multiplication assays. **a)** Sexual conversion rates of 3D7 wild type (WT), JmjC1-KO and JmjC1/2-dKO parasites cultured in minimal fatty acid medium supplemented with 2 mM choline chloride (+choline) (baseline sexual commitment rates) (open bars) or not (-choline) (sexual commitment-inducing conditions) (filled bars) in three replicates as assessed parasite counting using flow cytometry after 5 days of GlcNAc treatment to kill off asexual parasites (see methods section). Values represent the mean of three independent experiments. Dots represent individual values and error bars define the standard deviation. Significance was calculated using a one-way Anova t-test. p-value is shown as * < 0.05. **b)** Proportion of Pfg377+ (red) and Pfg377-(blue) cells as assessed by IFAs on Day9 gametocytes. Female gametocyte ratios (Pf377+) are displayed on the right side. Total numbers of gametocytes counted for both 3D7/JmjC1-KO and 3D7/JmjC1/2-dKO are low due to low sexual conversion rates. **c)** Representative flow cytometry plots of a 3D7 wild type (WT) culture (3D7 WT, TP2, replicate 1). Left: The plot shows the gate set to remove small debris (smaller than cell size), keeping the “cells” used for further gating. Middle: The “cells” population was gated to remove doublets, keeping “Single Cells” used for further gating. Right: SYBR green fluorescence intensity was used to separate uninfected from infected RBCs (iRBCs). The numbers indicate the percentage of cells included in the gate (i.e. the parasitemia).

**Supplementary Table S1.**
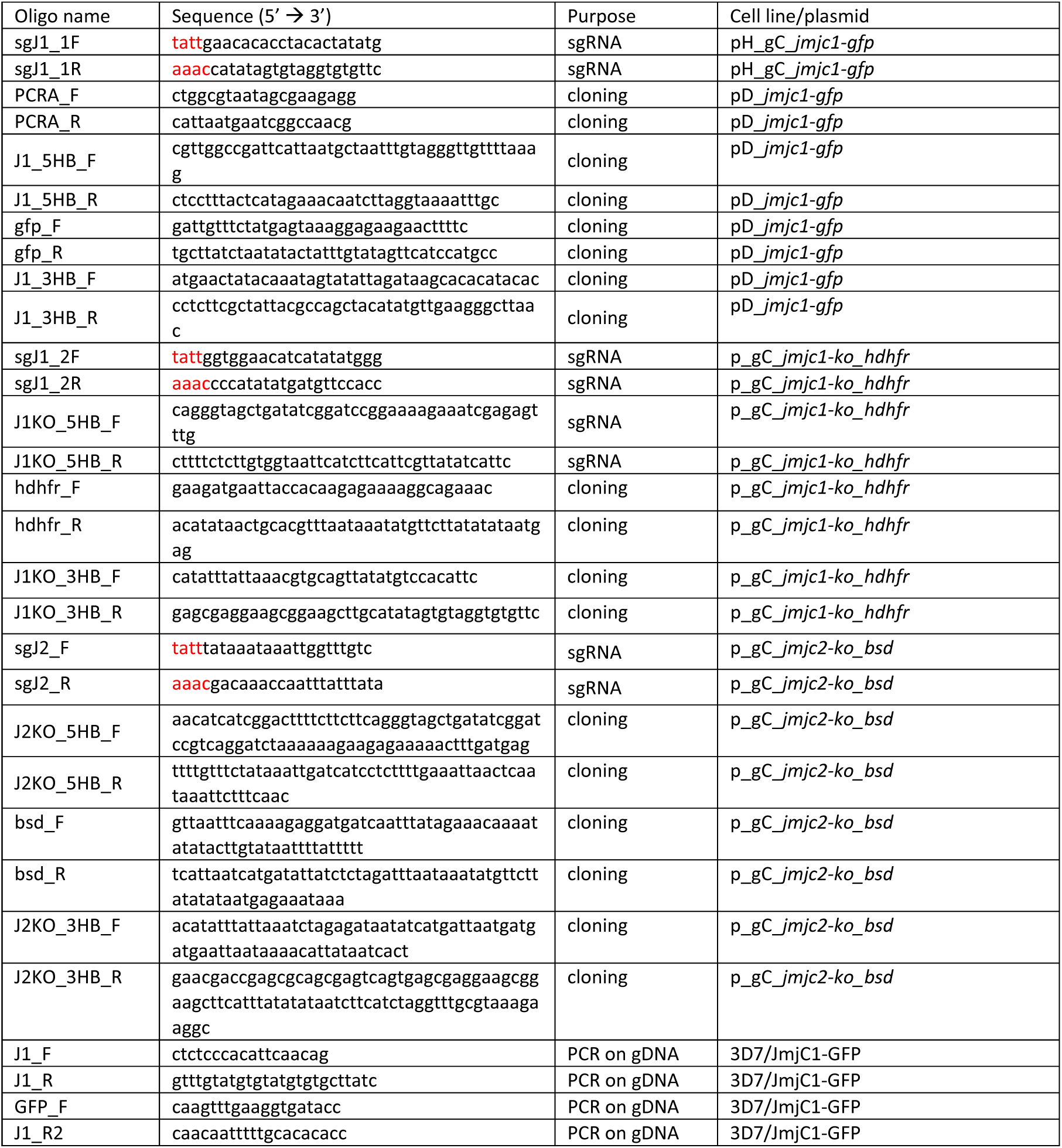
Oligonucleotides used in this study. Note: compatible single-stranded overhangs are highlighted in red.

